# Integrating Load-Cell Lysimetry and Machine Learning for Prediction of Daily Plant Transpiration

**DOI:** 10.1101/2024.11.24.625038

**Authors:** Shani Friedman, Nir Averbuch, Tifferet Nevo, Menachem Moshelion

**Affiliations:** The Robert H. Smith Institute of Plant Sciences and Genetics in Agriculture, The Robert H. Smith Faculty of Agriculture, Food and Environment, The Hebrew University of Jerusalem, Rehovot 7610001, Israel

**Keywords:** Artificial Intelligence, Feature Importance, Gravimetric Lysimeters, Irrigation Management, Machine Learning (ML), Precision Agriculture, Penman-Monteith, Transpiration-prediction

## Abstract

- We conducted research to predict daily transpiration in crops by utilizing a combination of machine learning (ML) models combined with extensive transpiration data from gravimetric load cells and ambient sensors. Our aim was to improve the accuracy of transpiration estimates.
- Data were collected from hundreds of plant specimens growing in two semi-controlled greenhouses over seven years, automatically measuring key physiological traits (serves as our ground truth data) and meteorological variables with high temporal resolution and accuracy. We trained Decision tree, Random Forest, XGBoost, and Neural Network models on this dataset to predict daily transpiration.
- The Random Forest and XGBoost models demonstrated high accuracy in predicting the whole plant transpiration, with R² values of 0.89 on the test set (cross-validation) and R^2^ = 0.82 on holdout experiments. Ambient temperature was identified as the most influential environmental factors affecting transpiration.
- Our results emphasize the potential of ML for precise water management in agriculture, and simplify some of the complex and dynamic environmental forces that shape transpiration.

## 1. Introduction

Transpiration, the process of water evaporation from plants, has been a subject of scientific studies for centuries. It primarily occurs through leaves, facilitating a continuous transport of water and essential nutrients from the roots. One remarkable aspect of transpiration is how plants regulate it in response to their surroundings through dynamic control mechanisms. This process is intricately linked to the behavior of stomata, small pores found on leaf surface. Stomata can adjust their aperture to manage the rate of transpiration (Lange et al., 1971). This adaptive response to the environment appears to have evolved as one of the protective mechanisms against excessive dehydration and physiological damage (Iqbal et al., 2020).

The dynamic ability of plants to regulate their transpiration rates plays a vital role in facilitating the exchange of carbon dioxide and water, thereby improving water use efficiency and optimizing growth. Various factors, both biological, such as plant size (Geller & Smith, 1982) and environmental such as solar Radiation (Pieruschka et al., 2010), temperature (Ben-Asher et al., 2008), humidity (Rawson et al., 1977), soil water supply (Madhu & Hatfield, 2014), carbon dioxide levels (Imai & Murata, 1976; Madhu & Hatfield, 2014), and wind speed (Dixon & Grace, 1984), can impact the transpiration rates of different plants. The understanding of these influential factors holds paramount significance in the fields of agriculture and ecophysiological sciences.

Understanding and quantifying the factors which affects transpiration and plant-water relations has a key role in optimizing water management strategies in agricultural and greenhouse settings. Therefore, various modeling approaches have been employed to simulate and evaluate transpiration rates in plants. One common approach is the use of mechanistic models, which are based on physiological principles and the understanding of plant structure and function. For example, the semi-empirical Ball-Berry model integrates stomatal conductance and environmental variables to estimate transpiration rates (Ball et al., 1987). However, it’s important to note that the Ball-Berry model simplifies the complex process, assuming a linear relationship between stomatal conductance and photosynthesis, which can significantly influence stomatal behavior and transpiration rates.

In contrast, process-based models like the Penman-Monteith equation take a more comprehensive approach. Process-based models represent a modeling approach that simulates natural processes by representing the underlying mechanisms. Penman-Monteith equation combine energy balance and aerodynamic principles to predict evapotranspiration (Monteith, 1965; Penman, 1948). The FAO56 Penman–Monteith is commonly used in the agriculture community to estimate evapotranspiration (Landeras et al., 2008). However, previous studies have found daily FAO56 Penman–Monteith exhibited up to 22% discrepancy in accuracy compared to the actual whole-plant transpiration measured using lysimeters (Averbuch & Moshelion, 2024; Kiraga et al., 2023; López-Urrea et al., 2006). These discrepancies underscore the ongoing need for improvement and refinement in modeling techniques to better understand and predict transpiration in different conditions and environments.

An alternative method involves developing models through machine learning algorithms, including artificial neural networks (ANN) and support vector regression (SVR). These have shown promising results in estimating transpiration based on environmental inputs and plant characteristics (Balasubramanian & Thirugnanam, 2023; de Meneses et al., 2020; Ferreira et al., 2019; Xing et al., 2022). However, many of these studies rely on broad or indirect data labels (e.g., remote sensing, estimated transpiration, or synthetic data) that do not fully capture the detailed plant–environment interactions critical for accurate physiological modeling. To extend our understanding of how machine learning has been applied in this field, we conducted a systematic search of the Scopus database for relevant studies (for more details, see Supplementary Methods S1). Among these, 33.9% (184 articles) employed indirect estimation methods, primarily remote sensing (150 articles) and eddy covariance (34 articles), while empirical models like FAO-56 Penman–Monteith appeared in 26.0% (141 articles). However, only 4.8% (26 articles) included keywords related to direct physiological measurements, such as sap flow sensors, load cells, lysimeters, porometry, or gas exchange systems—highlighting a clear gap in capturing direct plant-environment interactions.

Despite growing interest in ML-based models for transpiration prediction, accurately modeling whole-plant transpiration remains a major challenge. This is because transpiration is a highly dynamic process, regulated by thousands of signalling and transport processes within the plant and influenced by multiple, interdependent environmental variables. To capture this complexity, ML models must be trained on reliable, high-resolution physiological ground truth data that reflect actual plant behavior-not estimated or proxy data streams. Load-cells lysimeters are widely regarded as the gold standard for measuring whole-plant transpiration, as they enable direct quantification of evapotranspiration (ET) or transpiration flux (Halperin et al., 2017). ML models trained on data derived from lysimeters can serve as reliable benchmarks for validating or calibrating alternative, indirect estimation methods (Amani & Shafizadeh-Moghadam, 2023; Anapalli et al., 2016; X. Liu et al., 2017). Continuous, high-throughput whole-plant transpiration measurements from load-cells lysimeters provide the necessary physiological precision for this task—they represent the direct, non-manipulated, transpiration ‘ground truth’ required to train accurate and biologically meaningful models. Without such direct measurements, models risk learning patterns that do not reflect real plant behavior, ultimately limiting their accuracy, robustness, and utility. Despite their benefits, most studies employing lysimeters tend to use either drainage or manual lysimeters, resulting in a limited number of instruments due to the high costs and labor involved in deployment and maintenance (Z. Chen et al., 2020; Kiraga et al., 2023). Our approach, by contrast, leverages a high-throughput load-cells lysimeter platform, with high signal to noise ratio, enabling large-scale, directly labeled datasets suitable for robust model training.

In this study, we focused on daily measurements of transpiration for mediterranean summer and winter crops, tomato and cereals (Wheat and Barley), respectively. Over the past seven years, we have collected a precise-data set of direct physiological traits from hundreds of plant specimens, along with corresponding environmental data. Leveraging this unique dataset, we employed machine learning models to predict the daily transpiration of well-irrigated plants. Our data is distinguished by its high scale and the use of ground truth annotated physiological measurements (e.g., whole plant transpiration g/day). These are obtained from an extensive array of load cells lysimeters and atmosphere conditions.

The objectives of this study are to leverage our dataset, which includes soil-plant-atmosphere continuum data, to (1) evaluate the effectiveness of conventional tree-based machine learning models, including Random Forest, a XGBoost model, and a neural network model, in predicting daily transpiration for tomato and cereal crops; and (2) determine the hierarchical significance of diverse ambient factors, thereby pinpointing the critical environmental factors that exert influence on transpiration.

## 2. Materials and methods

### 2.1 Research overview

In this study, we aimed to understand the determinants of daily transpiration in plants using a structured machine learning research workflow. Data collection took place within semi-commercial greenhouses using functional phenotyping platform composed of load-cells lysimeters. This platform provided essential information on several plant physiological traits (transpiration, plant weight, growth rate etc.) and ambient conditions After essential data pre-processing steps, including outlier removal and transformation to daily values, we had 6115 observations (81% of the original daily data), split into training and testing sets. Machine learning models, including tree-based models and neural network, were trained to predict daily transpiration using the features: plant weight, temperature, humidity, Daily Light Integral (DLI), Vapor Pressure Deficit (VPD), plant type, and soil type. Model evaluation encompassed various metrics such as R^2^ and Root Mean Square Error (RMSE). Model interpretation was accomplished through the analysis of feature importance using techniques like Permutation and Shapley Additive Explanations (SHAP; see 2.9). These methods hold major contribution in understanding the different features to the models’ predictions and their impact on daily transpiration. This workflow allowed us to gain valuable insights into the determinants of daily transpiration in plants.

### 2.2 Experimental site

Data was collected from two semi-controled greenhouses located at the I-CORE centre for Functional Phenotyping of the Faculty of Agriculture, Food, and Environment in Rehovot, Israel (http://departments.agri.huji.ac.il/plantscience/icore.php). The ‘main greenhouse’ (Fig. 1A) is a polycarbonate covered structure at coordinates 31°54’15.0”N, 34°48’03.6”E, measuring 18 m in length and 16 m in width, with a gutter height of 4.5 m and a maximum ridge height of 6 m (dual-gable structure). The estimated internal volume, based on an average height of 5.25 m, is approximately 1,417.5 m³. The ‘secondary greenhouse’ (Fig. 1B), also polycarbonate covered, located at 31°54’22.5”N, 34°48’10.6”E, spans 6 x 17 m^2 meters and reaches a height of 3 m.

**Figure 1:**
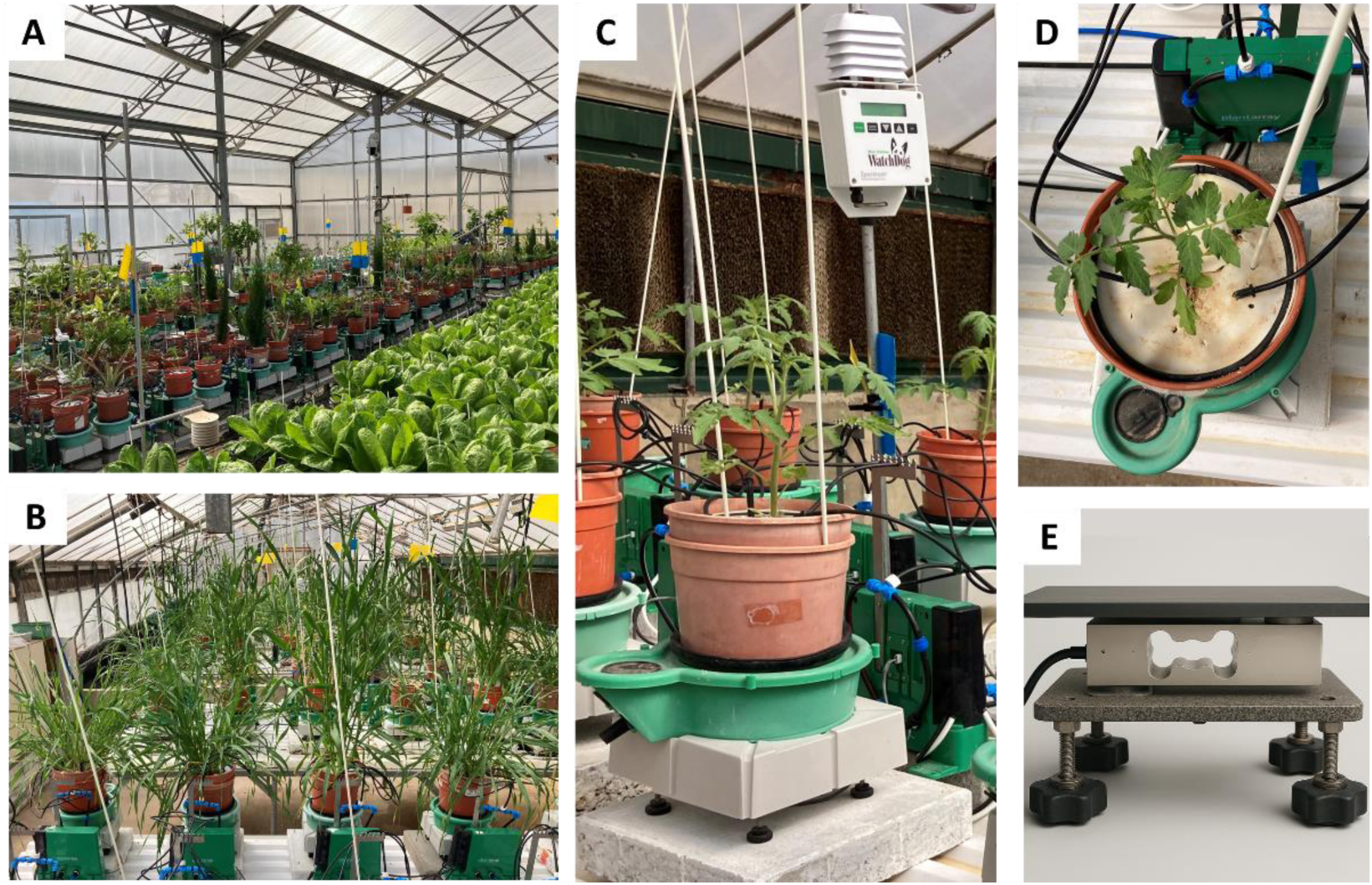
Configuration of load cell lysimeter system and greenhouse. (A) Main greenhouse containing various crops monitored using the PlantArray phenotyping platform. (B) Secondary greenhouse with wheat plants monitored using the same system. (C) Image of a single PlantArray unit, showing the load-cell lysimeter, double-pot setup, “personalize” controller collecting data and controlling each pot irrigation, and the central weather station used for continuous environmental monitoring. (D) Top view of a pot showing the sealed soil surface cover, designed to isolate plant transpiration from soil evaporation. (E) Photorealistic illustration of a single lysimeter unit. It includes a temperature-compensated load cell that converts mechanical force into an electrical signal, directly connected to the controler. The cell is mounted between two steel platforms to ensure stable weight measurement. As shown in (C), the lysimeter is typically covered with a polystyrene block and a thermal-isolated lid.

Both facilities feature natural daylight conditions and are equipped with a cooling pad along the northern wall to maintain temperatures below 35 degrees Celsius. Ventilation is achieved by using a positive pressure cooling system, where outside air is pushed into the greenhouse through a wet pad by four fans, each with capacity of 18,000 m^3^/h, totaling flow rate of 72,000 m^3^/h (This setup enables an air exchange rate of approximately 51 air changes per hour, for the main greenhouse). The fans are automatically activated about 30 minutes before sunrise and turned of about 30 minutes after sunset to ensure consistent airflow during the photoperiod. This high ventilation rate is designed to maintain CO₂ concentrations similar to those in the ambient outdoor air, facilitating natural plant responses. To complement these efforts and further simulate natural external conditions, wind speed and direction are precisely monitored using an ATMOS 22 ultrasonic anemometer, centrally located in the main greenhouse. This sensor records an average wind speed of 0.324 m/s. For both greenhouses, the experimental conditions varied, with natural Daily Light Integral (DLI) values ranging from 2 to 34 mol/(m²·d) and mean daily temperatures fluctuating between 10 and 33°C (Table S1A).

### 2.3 General experiment setup and Data collecting

The data was collected from June 2018 to March 2024. Using the functional phenotyping platform – PlantArray (PlantDitech, Israel; Fig. 1C) as described by Dalal et al., 2020; Halperin et al., 2017. Briefly, an array of load cell lysimeters was utilized (Fig. 1), to continuous plant weight measurement and the derivation of both transpiration-induced water loss and daily plant mass accumulation. To ensure that water loss reflects plant transpiration only, the surface of the soil in each pot was sealed to prevent soil evaporation (Fig. 1D). All data was automatically collected and uploaded to the cloud base SPAC analytics system (PlantDitech, Israel).

Meteorological data, consisting of four essential variables - temperature, VPD, light, and relative humidity (RH), were collected using a weather station (Watchdog 2000 series; Spectrum Technologies, Illinois, USA) connected to the PlantArray system. Within each greenhouse, the atmospheric sensors were positioned at the center of the greenhouse, approximately 1 meter above the height of the pot surface (Fig. 1C).

The lysimeter-based phenotyping system employed several strategies to enhance the signal-to-noise ratio, thereby reducing potential artifacts in the noisy greenhouse environment. These strategies include the use of high-accuracy load cell transducers, achieving a precision of ±0.167 g per kg loaded on each cell. These transducers are also temperature-compensated to effectively minimize signal drift caused by ambient temperature fluctuations. Furthermore, each load cell is connected via a short cable (45 cm) to its individual analog-to-digital (A/D) controller, significantly reducing analog electrical interference and noise (typically associated with long cables connected to a single data logger). To prevent overheating due to direct solar radiation, thermal insulation and sealed covers are applied separately to each load cell. Additionally, vibration-induced noise is mitigated by placing compressed foam cushions and mass under each load cell. Measures to counteract the “pot effect” (Gosa et al., 2019), such as double-pot arrangements isolating the soil and roots from direct solar radiation induced heat, to ensure the reliability and consistency of the physiological data collected during the experiment. (Fig. 1E)

#### 2.3.1 Plant Material and Growing Media

The plants in the greenhouse were grown in pots (4L) filled with sand (Silica sand grade 20-30, particle size 0.595–0.841 mm; Negev Industrial Minerals Ltd., Yeruham, Israel) or soil (Bentel 11 garden mix, composed of (w/w) 55% peat, 20% tuff and 25% puffed coconut coir fiber; Tuff-Substrates, Alon Tavor, Israel).

Tomato, wheat and barley (cereals), were employed in this experiment. The tomato (*Solanum lycopersicum*) variety incorporated in the study was mainly M82 cultivar. The cereals included *T. turgidum subsp. durum ‘Svevo’* and *T. aestivum cv. Gadish*, among others. Focusing on these commonly cultivated crops, ensuring multiple repetitions, and capturing the diversity of both winter and summer representations.

#### 2.3.2 Irrigation and water balance measurements

All pots were irrigated through repeated irrigation and drainage cycles every night, reliably restoring the soil to field capacity (hereafter referred to as “well-irrigated”). This approach not only maintained optimal soil moisture levels but also facilitated leaching to prevent salt accumulation. Daily pre-dawn pot mass was measured after full drainage. Plant mass was calculated at this point. The difference in pot mass between consecutive days reflects the plant’s biomass gain. The lack of additional irrigation throughout the daylight hours ensured a monotonic pot-mass decrease between subsequent irrigation events. Transpiration was calculated based on the mass loss during the day. For full details on calculations and system see Dalal et al. (2020) and Halperin et al. (2017).

In this study, we focused exclusively on non-stressed plants by ensuring that soil water content consistently remained above the transpiration-limiting threshold (Halperin et. al., 2016). This was achieved by selecting data exclusively from control (non-drought) treatments, while deliberately excluding any data from plants that surpassed their pot’s capacity, thereby ensuring non-pot effect of draught conditions. Therefore, we checked the data manually, removing entries where plants exhibited ‘pot limitation’, a scenario where a plant’s transpiration plateaus due to reaching the pot’s maximum available water capacity. While the calculated total water-holding capacity is approximately 860 g in sand and 1700 g in potting media (Dalal et al., 2020), the actual available water for plants is lower: around 600 g/day in sand and 1200 g/day in soil, influenced by root structure and soil properties.

### 2.4 Data pre-processing

The data was initially collected at 3-minute intervals. For this research, it was further processed to generate daily values, covering the period from first light to last light (Table 1). Using daily values reduces the total data volume to balance capturing environmental variability while keeping data processing manageable. Moreover, this simplifies data interpretation, making it practical for applications such as irrigation decisions, much like other common methods, including the FAO56 Penman–Monteith model for irrigation management.

**Table 1:** Dataset description and the main preprocessing methods.

| category | variable | Units | Description |
| --- | --- | --- | --- |
| ID | Timestamp |  | Date |
|  | lcore greenhouse |  | The greenhouse the data was collected from. Encoded as Main = 1 and secondary = 0. |
|  | expld |  | Experiment number: each independent experiment contained 1-10 plants (repetitions) that the data was collected simultaneously from. A total of 48 independent experiments were used. |
|  | plantId |  | The plant specific ID number, each plant was measured continuously for about 3-4 weeks. On average 23 daily observations per-plant. A total of 269 plants were used. |
| Meteorology | Temperture | °C | The average temperature at daytime (from the first to last light). |
|  | RH | % | The average Relative Humidity at daytime (from the first to last light). |
|  | VPD | kPa | The average Vapor Pressure Deficit (VPD) at daytime (from the first to last light). |
|  | DLI | mol/m<sup>2</sup>*day | Daily Light Integral (DLI) was calculated using formula 1. |
| Others | Potting media |  | “Encoded soil” - dummy variables, 0 =silica sand, 1 = soil (4508 and 1607 observation respectively). |
|  | Plant types |  | “Encoded plant” - dummy variable, 0 =tomato, 1 =cereal (3975 and 2140 observation respectively) |
| Plant weight | Plant Weight | g | The net weight of the plant as calculated at 4AM, after soil field capacity (for details see Halperin et al 2017). |
|  | Transpiration | g | Daily water evaporating trough the plant (for details see Halperin et al 2017). This variable was used as output for the models. |

The type of potting media (sand or soil) and the specific plant species (tomato or cereal) were recorded. Light data in terms of Photosynthetically Active Radiation (PAR) was measured at 3-minute intervals. The Daily Light Integral (DLI) was calculated using the formula:

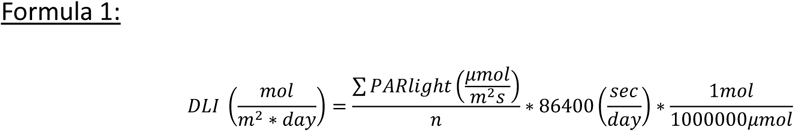

- *n* - number of samples in a full day (480 if sampled every 3 min).
- 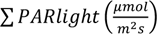 is the sum of PAR light measurements throughout the day.

This calculation provided insights into the cumulative light exposure experienced by the plants over the course of a day.

Our dataset initially included 7,547 observations, where each observation representing a single plant measured on a given day. An example of an individual plant is illustrated in Fig. 2, with a single observation shown in Fig. 2H. After a comprehensive data cleaning process, we concluded with a total of 6,115 observations. This reduction was due to the removal of observations with missing values, extreme outliers, and those exhibiting irregular behaviours such as weight loss, water shortage, or illnesses. The final variables and some pre-processing methods, such as daily aggregation of values and encoding, are detailed in Table 1.

**Figure 2:**
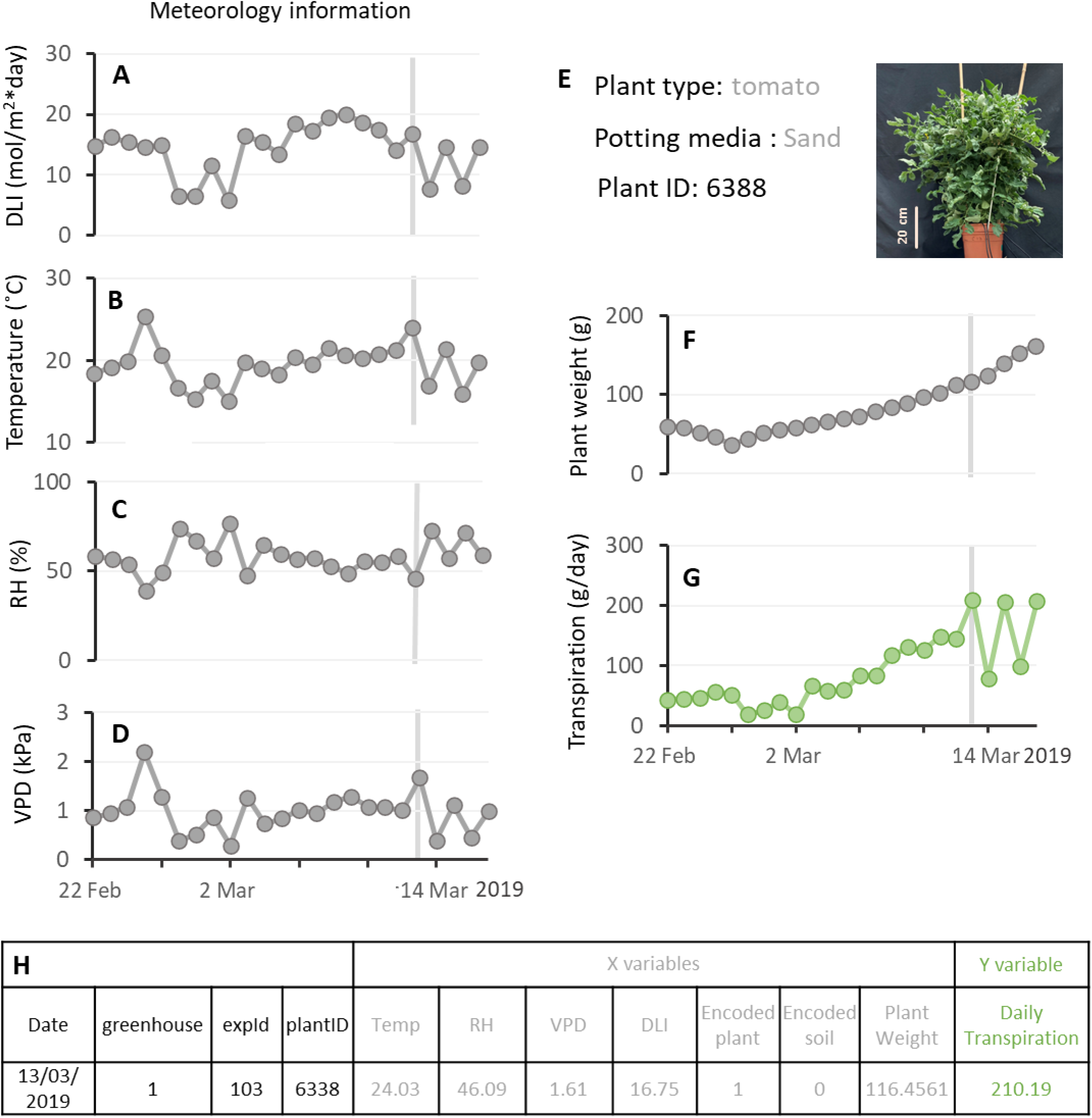
Representative meteorological and physiological data derived from the I-core greenhouse PlantArray system. This figure presents data from a single plant as an example. Daily average of (A) light integral, (B) temperature, (C) Relative humidity, and (D) vapor pressor deficit over a period of 22 days. (E) Plant identity information: plant type, potting media, ID number and plant picture at the end of the experiment. (F) Plant biomass - plant net weight (G) whole-plant daily transpiration, note the sensitivity of the transpiration to the methodological changes. (H) A single observation as it is documented it the data set. This observation is visually depicted in Figures A-F, on the 13th of March. X variables marked in gray and the y variable in green, representing the inputs and outputs used by computer models for learning and prediction.

### 2.5 Tuning, Training, and Testing Datasets

After preprocessing, the dataset comprised 6115 daily observations. Fig. 2 illustrates a sample plant with 24 observations, Fig. 3 presents the full dataset from the main greenhouse, including meteorological and aggregated physiological data. The feature variables included Temperature, Relative Humidity (RH), vapor pressure deficit (VPD), Daily light Integral (DLI), plant weight, plant type, and soil type, with the target variable being daily transpiration (Fig. 2H, Table 1).

**Figure 3:**
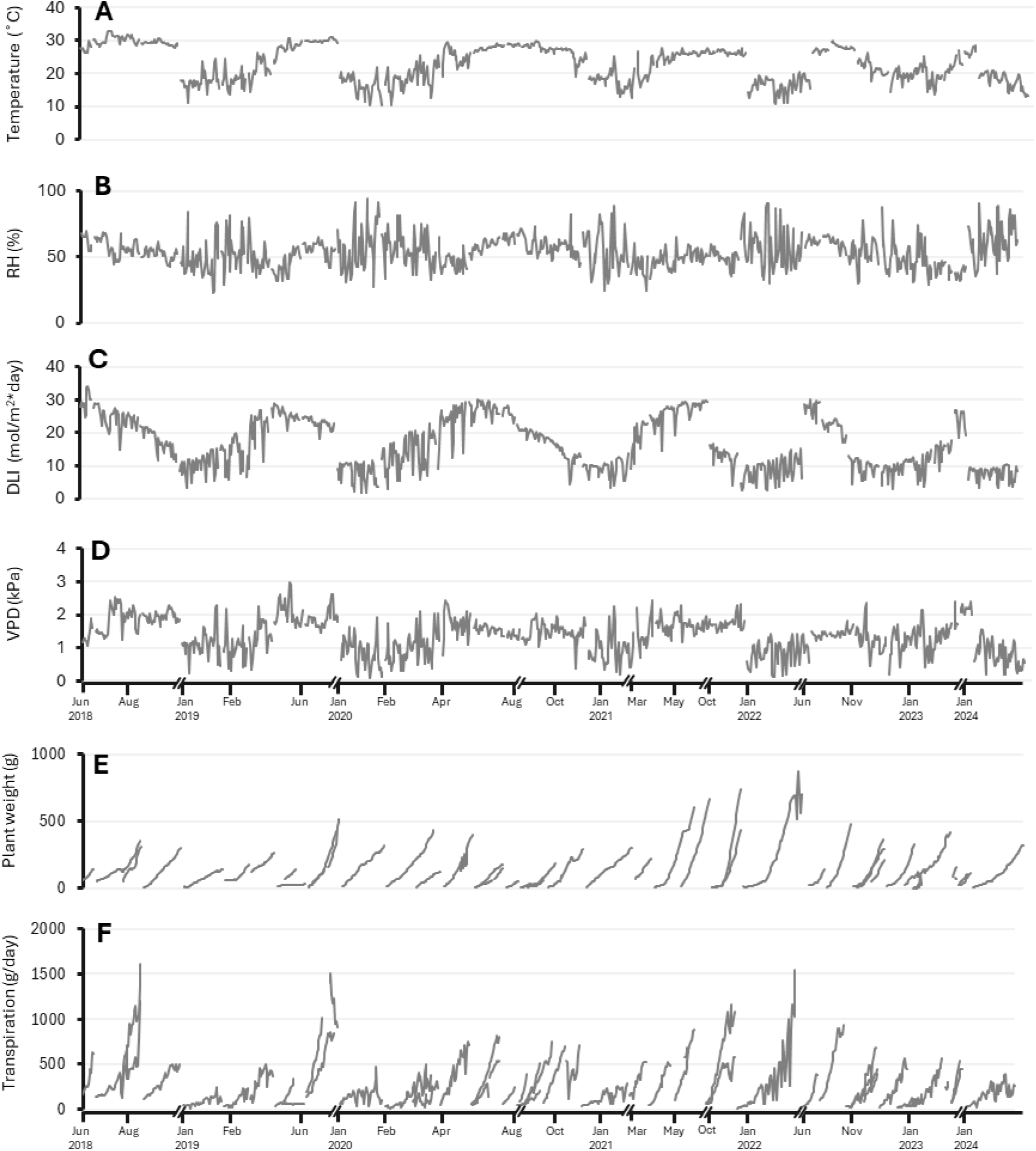
Summary of meteorological and physiological data collected in the main I-CORE greenhouse over the entire study period. This figure presents the full dataset collected from the PlantArray lysimeter system across multiple experiments conducted in the main greenhouse. Panels (A–D) show daily average meteorological conditions: (A) daily light integral (DLI), (B) temperature, (C) relative humidity (RH), and (D) vapor pressure deficit (VPD), recorded continuously throughout the experiments timeline. Panels (E–F) present physiological responses averaged across all plants measured in each experiment. (E) Average plant weight, illustrating the typical pattern of plant development, where plants begin small and progressively accumulate biomass over time. (F) Average daily transpiration per plant, reflecting physiological responses to both environmental conditions and developmental stage. Gaps along the x-axis correspond to time periods when no relevant experiments were conducted or when the greenhouse was inactive. Note that different experiments may overlap in time.

#### 2.5.1 Validation Dataset (Holdout)

To assess the model’s generalizability, a holdout dataset comprising 805 observations from 4 randomly selected experiments was established, a random seed function was set to ensure repeatability. This subset includes 400 observations related to tomato plants and 405 observations concerning cereal plants (for more information see Table S1b and Fig. S1b). This holdout dataset aims to reflect the model’s performance and effectiveness on unseen data.

#### 2.5.2 Training Dataset

Ninety percent (4779 observations) of the remaining data was randomly selected to tune the hyperparameters, as detailed in Section 2.7. This initial subset helped determining the optimal hyperparameters, allowing us to assess the effectiveness of the tuning process. After selecting the best hyperparameters, the models were then trained on the full dataset, excluding the holdout set (in total 5310 observations). This approach ensures that the final model training incorporates the broadest data spectrum available while retaining an independent holdout set to evaluate the model (see 2.9).

### 2.6 Machine learning models for estimating daily transpiration

#### 2.6.1 Decision Tree

Decision trees are a simple, yet powerful machine learning model used for classification and regression tasks. They recursively split the data into subsets based on the features to create a tree-like structure. At each node, the decision tree selects the feature and split point that minimizes impurity, aiming to create more homogenous subsets. The final predictions are made based on average value (for regression) of the samples in the leaf nodes. Decision trees are interpretable and can capture complex relationships in the data, but they may suffer from overfitting and lack generalization ability. The induction of decision trees is one of the oldest and most popular techniques for learning discriminatory models, which has been developed independently in the statistical (Breiman et al., 1984; Mingers, 1989) and machine learning (Quinlan, 1993) communities (Fürnkranz, 2011).

#### 2.6.2 Random Forest

Random Forest is an ensemble learning technique based on decision trees. It builds multiple decision trees, each trained on a random subset of the data and a random subset of the features. The final prediction is obtained by averaging (for regression) the predictions of individual trees. Random Forest overcomes the overfitting issue of decision trees and improves the model’s performance and robustness. It can handle high-dimensional datasets and capture interactions between features, making it a popular choice for various applications.

The random forest or decision tree forest is an algorithm created in 1995 by Ho (Ho, 1995), then formally proposed by scientists in 2001 (Breiman, 2001; Cutler & Zhao, 2001).

#### 2.6.1 XGBoost model

eXtreme Gradient Boosting (XGBoost) is a method, initially proposed by (T. Chen & Guestrin, 2016), that generally generates high accuracy and fast processing time while being computationally less costly and less complex. The XGBoost model is a scalable machine learning system for tree boosting. The XGBoost model integrates several “weak” learners for developing a “strong” learner through additive learning. Parallel computation is automatically implemented during training to enhance computational efficiency (Fan et al., 2021).

#### 2.6.3 Artificial neural networks

Neural networks are a class of deep learning models inspired by the structure and function of the human brain (Haykin, 2009; Rosenblatt, 1958). They consist of interconnected nodes (neurons) organized into layers. Each node applies an activation function to the weighted sum of its inputs to produce an output (Fukushima, 1969). As many ML models, Neural networks can learn complex and nonlinear relationships in the data through the process of forward and backward propagation during training (Leibniz, 1920).

### 2.7 Hyperparameters tuning

Hyperparameter tuning is the process of discovering the optimal configuration for the hyperparameters of a machine learning model to achieve the best performance. Hyperparameters are external configuration settings established before the training process and are not learned from the data. Examples included learning rates, regularization strengths, and the number of trees in a random forest.

This process of hyperparameter tuning was carried out through cross-validation grid search on the training data. Grid search cross-validation is a method employed to systematically explore a vast array of combinations of hyperparameter values. By leveraging 5-fold cross-validation, the dataset was divided into 5 subsets, allowing for multiple rounds of training and validation. This approach aimed to find the optimal set of hyperparameter values and ensure the model’s robustness by evaluating its performance across different subsets of the training data. The optimal hyperparameter values, and the options we tested are summarized in Table 2.

**Table 2:** Hyperparameters. Adjusting these parameters change the learning ability, so the model would not underfit neither overfit the dataset. Implementing GridSearch cross-validation on 90% of the training data set. Systematically all the possible combinations of the Hyperparameter options were evaluated and the optimal setting was retained. We used the 5-fold cross validation splitting strategy to test the optimal parameters on 5 different subsets of the training dataset.

| model | Hyperparameter | Explanation | Options | Optimal value | Combinations |
| --- | --- | --- | --- | --- | --- |
| Decision Tree | min_samples_split | The minimum number of samples required to split an internal node. | [2,3,4] | 4 | 60 candidates<br>* 5 folds = 300<br>fits |
|  | min_samples_leaf | The minimum number of samples required to be at a leaf node of the trees. | [1,2,3,4] | 4 |  |
|  | max_depth | The maximum depth of a decision tree. | [None, 5,7,10,12] | 10 |  |
| Random Forest | n_estimators | The number of decision trees (estimators) or boosting rounds in the random forest ensemble. | [50,60,70,80,100] | 100 | 240<br>candidates * 5<br>folds = 1200<br>fits |
|  | min_samples_split | Explained above | [2,3,4,5] | 4 |  |
|  | min_samples_leaf | Explained above | [1, 2] | 1 |  |
|  | max_depth | Explained above | [None, 12,15] | 15 |  |
|  | bootstrap | This hyperparameter determines whether bootstrap samples (sampling with replacement) are used when building trees in the random forest. | [True, False] | True |  |
| XGBoost | n_estimators | Explained above | [60, 80, 100] | 100 | 1296<br>candidates*5<br>folds = 6,480<br>fits |
|  | min_child_weight | Minimum sum of sample weight (hessian) of the smallest leaf node. Larger values, preventing splits that have a smaller hessian value. | [1, 2, 4, 5] | 1 |  |
|  | max_depth | Explained above | [3, 5, 6, 7] | 7 |  |
|  | gamma | Minimum loss reduction required to make a further partition on a leaf node of the tree (helps in pruning the tree). | [0, 0.1, 0.2] | 0 |  |
|  | reg_lambda | L2 regularization term on weights. It adds a penalty to the loss function based on the size of the weights. | [0, 1, 2] | 0 |  |
|  | learning_rate | scales the contribution of each tree in the ensemble. A smaller learning rate makes the algorithm more robust by reducing the step size. | [0.3, 0.1, 0.01] | 0.1 |  |
| Neural Network | num_units – layer 1 | controls the number of units (neurons) in each hidden layer of the neural network. It represents the width of the network. | [32, 64, 96, 128, 160, 192, 224, 256, 288] | 32 | 1296 candidates *<br>40 epoch =<br>51,840 fits |
|  | num_units – layer 2 | Explained above | [32, 64, 96, 128, 160, 192, 224, 256, 288] | 288 |  |
|  | optimizer | determines the optimization algorithm used to update the model's weights during training. The code considers two options: 'adam' (Adaptive Moment Estimation) and 'rmsprop' (Root Mean Square Propagation) | ['adam', 'rmsprop'] | rmsprop |  |
|  | Activation - Layer1 | The activation function applied to the output of each neuron in the hidden layers. Common choices are 'relu' (Rectified Linear Unit) and 'tanh' (Hyperbolic Tangent). Activation functions introduce non-linearity to the network, enabling it to learn complex patterns. | ['relu', 'tanh'] | tanh |  |
|  | Activation- Layer 2 | Explained above | ['relu', 'tanh'] | tanh |  |
|  | learning_rate | The learning rate controls the step size at which the optimizer updates the model's weights in each iteration during training. It is a crucial hyperparameter as it affects the speed and stability of training. | [0.001, 0.01] | 0.01 |  |

### 2.8 Modelling process

We implemented our models using a robust set of Python libraries tailored for statistical computing, data manipulation, and machine learning. Specifically, we used: Scipy (Virtanen et al., 2020), for conducting statistical tests, Numpy (Harris et al., 2020), for high-performance numerical operations, Pandas (Mckinney, 2010; The pandas development team, 2020), for data handling and manipulation, Sklearn (Pedregosa et al., 2011) for machine learning algorithms and model evaluation tools, Keras (Chollet, 2015) and TensorFlow (Abadi et al., 2016) for building and training neural network models.

### 2.9 Cross-Validation and Performance Evaluation

Cross-validation was conducted on the dataset to validate model stability and reliability. We used 10-fold cross-validation, allowing each subset of data to be used as both training and testing sets iteratively, ensuring comprehensive performance assessment. This method not only helps in assessing the performance across different slices of data but also in comparing the effectiveness of different models under varying conditions (Table 3).

**Table 3:**
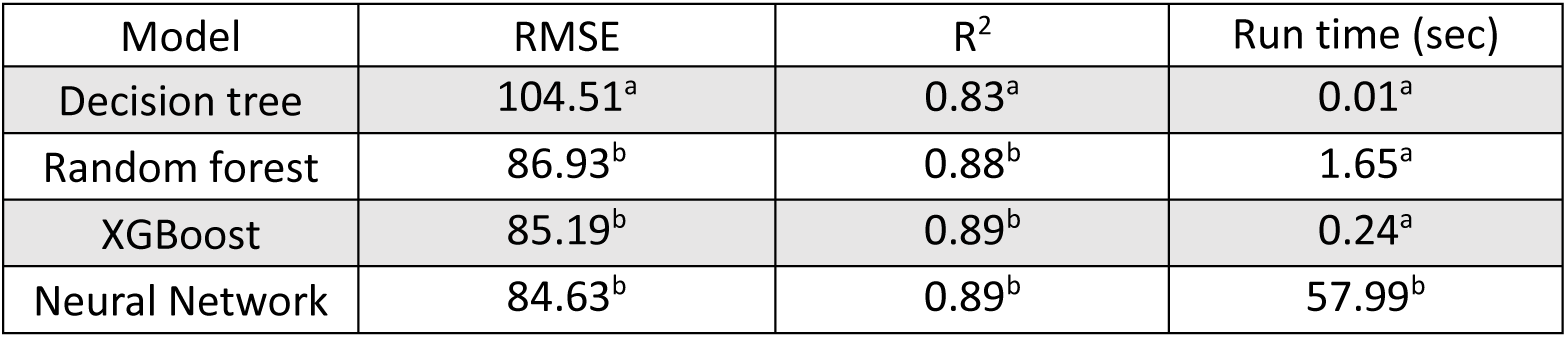
Comparison of evaluation scores among the four methods models. The average of 10-fold Cross validations are presented. An ANOVA test was conducted to statistically evaluate performance differences among the 10-fold models scores, followed by Tukey’s Honest Significant Difference (HSD), differences are indicated by letters.

The final model evaluation was performed on the holdout set, using metrics such as Root Mean Square Error (RMSE), Mean Absolute Error (MAE), and the R² statistic to compare the predictions and actual measurements (Table 4). These metrics provided a detailed measure of model accuracy, prediction error, and the proportion of variance explained by the model.

**Table 4:** model evaluation statistics on the holdout experiments data.

| Model | R<sup>2</sup> | RMSE (g) | MAE | Relative error<sup>1</sup> (%) | Residual<sup>2</sup> (g) | Residual<sup>2</sup> (%) | Tukey HSD<sup>3</sup> |
| --- | --- | --- | --- | --- | --- | --- | --- |
| Decision Tree | 0.77 | 98.65 | 75.53 | 64.71 | 75.53 | 64.71 | a |
| Random Forest | 0.81 | 90.77 | 69.71 | 60.12 | 69.71 | 60.12 | a |
| XGBoost | 0.82 | 88.41 | 66.71 | 57.37 | 66.71 | 57.37 | a |
| Neural Network | 0.74 | 106.00 | 79.11 | 64.60 | 79.11 | 64.60 | a |

<sup>1</sup> Relative error, also known as the Relative RMSE, is a measure that quantifies the accuracy of predictions relative to the scale of the actual values. is calculated as the RMSE divided by the mean of the actual values
<sup>2</sup> The mean absolute residual was calculated.
<sup>3</sup> Tukey test on the residuals between the predicted and actual values. Different letters indicate significant difference in Tukey HSD test $P_v < 0.05$ .

### 2.10 Model Validation Using External Greenhouse and Growth Room Data

To further evaluate the generalizability of our machine learning models, we tested their performance on two external datasets collected independently from the training and validation data. The first dataset was obtained from a greenhouse facility located in Tel Aviv University, operated entirely by a different research group, using separate personnel, experimental planning, and management protocols. Although this facility employed the same phenotyping platform (PlantArray; PlantDitech, Israel) and followed a similar measurement protocol, the experiments were conducted independently from those at the I-CORE center in Rehovot. Data were collected using the SPAC analytics system and included plant weight and transpiration measurements from experiments carried out between April 17, 2024, and November 16, 2024.

For external testing, we selected three individual plants, each from a different experiment. Since the Tel Aviv team includes soil bulk weight in their lysimeter measurements, we standardized the datasets by adjusting the initial seedling weight to 10 g to match the conventions used in our own experiments, where only net plant biomass is recorded. This adjustment ensured consistency and comparability between the datasets.

The second external dataset was collected from our indoor growth room (Room 101; Fig. S2), located within the I-CORE center, but operated under different environmental conditions and experimental constraints compared to the main greenhouse. This dataset offers an independent context due to the controlled lighting, humidity, and temperature conditions specific to growth room setups.

Both external datasets were evaluated using our Model Testing App, a web-based interface designed to assess model performance on new user-provided datasets. Users with SPAC analytics access can upload their experimental data and receive transpiration predictions generated by our pre-trained models, along with performance metrics for direct comparison against measured values. The application is publicly available at: https://test-daily-transpiration-model-spac-user.streamlit.app/.

### 2.11 feature importance

To interpret the predictions and understand the importance and contribution of each feature to building the models several feature important tests were presented:

#### 2.11.1 Impurity-based feature importance

Impurity-based feature importance is a technique commonly employed in decision tree-based machine learning models, to assess the significance of individual features in making predictions. The method calculates the contribution of each feature by measuring how much it reduces the impurity (e.g., Gini impurity or entropy) in the model’s decision nodes. This approach is simple, computational efficient, and interpretable, as it provides a clear ranking of features based on their impact. However, Impurity-based feature importance tends to favor variables with more categories or levels, which can lead to biases.

#### 2.11.2 Permutation feature importance

Permutation feature importance is a technique used to evaluate the significance of individual features in a machine learning model. The process involves systematically shuffling the values of a single feature in the dataset and observing the impact on the model’s performance. By comparing the model’s performance before and after the permutation, a decrease in performance indicates that the feature is crucial, as its alteration disrupts the model’s predictive accuracy. This approach does not require retraining the model and can be applied to various algorithms (not just tree-based models). However, it may not capture feature interactions and is most effective when dealing with uncorrelated features. In our study, the “sklearn.inspection.permutation_importance” was used.

#### 2.11.3 SHapley Additive exPlanations (SHAP)

SHapley Additive exPlanations (SHAP) by Lundberg et al., 2017 is a method to explain individual predictions. SHAP is based on the game theoretically optimal Shapley values (Molnar, 2022). Shapley values provide a mathematically fair and unique method to attribute the payoff of a cooperative game to the players of the game (Merrick & Taly, 2020; Shapley, 1953). SHAP values help us understand the role of each feature in a prediction by calculating how much each feature has pushed the prediction higher or lower, compared to prediction without that feature. For instance, applying SHAP to our model (Fig. S4 and see Fig. 6) revealed that temperature significantly influences predicted daily transpiration. Lower temperatures (depicted in blue) correlate with negative SHAP values, pushing predictions downward, while higher temperatures (depicted in red) associate with positive SHAP values, pushing predictions upward. Therefore, in this context, the model suggests that transpiration tends to be lower at lower temperatures. The SHAP library in python was used (Lundberg et al., 2017). For more info see https://shap.readthedocs.io/en/latest/example_notebooks/api_examples/plots/beeswarm.html#

### 2.12 Statistical Analysis

Several statistical methods were employed in this study. To assess the correlation between features, we used the correlation coefficient (r), which measures the strength and direction of the linear relationship between two variables. For evaluating model performance, we used R² (coefficient of determination), which represents the proportion of variance in the dependent variable that is predictable from the independent variables. T-tests were used to compare the means of two groups and to assess whether there were significant differences between them. P-values (Pv) associated with the t-tests were used to determine statistical significance, with a threshold of Pv < 0.05 considered significant.

For categorical variables, we used a chi-square test to evaluate associations between categories. The chi-square value indicates the magnitude of the discrepancy between observed and expected values, while the p-value reveals whether this discrepancy is statistically significant. To compare multiple groups and determine whether their means differed significantly, we applied ANOVA (Analysis of Variance), calculating the F-statistic and corresponding Pv to evaluate overall model significance. Additionally, when group variances were unequal, we used Welch’s test as a robust alternative to ANOVA, reporting the F-ratio and Pv to verify the statistical significance of the differences.

## 3. Results

From June 2018 to March 2024, multiple experiments were conducted in our greenhouses using the PlantArray systems. These experiments yielded continuous data on both plant physiology and environmental conditions. We collected, cleaned, labeled, and averaged the data to obtain daily values, as described in the methods section 2.4. For this research, we have used only well irrigated data (soil water content at the end of the day was higher than the total daily transpiration; see methods 2.3.2), resulting in a dataset comprising 6115 observations, with each observation representing a single plant, and its ambient conditions in a day of measurement.

This article investigates the environmental responses of tomato and cereal crops, cultivated during summer and winter seasons respectively. Our findings reveal significant differences in environmental conditions between the two (Fig. 4A and Fig. S3). Tomato plants were exposed to significantly higher summer temperatures (grand mean 25.3°C for tomatoes and 18.6°C for cereals, p < 0.001; Fig. 4A1) and relative humidity (grand mean 56.15% for tomatoes and 52.49% for cereals, p < 0.001; Fig. 4A2) compared to the winter conditions experienced by the cereals. Consequently, tomatoes plants were exposed to higher daily average vapor pressure deficit (VPD) than cereals (grand mean 1.53 kPa and 1.22 Kpa respectively; Pv<0.001; Fig. 4A3). The Daily Light Integral (DLI) for tomatoes was also greater (Pv<0.001), with an average of 18.27 mol/m²/day and a peak of 33.99 mol/m²/day, whereas cereals recorded a lower average DLI of 11.36 mol/m²/day and a maximum of 28.96 mol/m²/day (Fig. 4A4).

**Figure 4:**
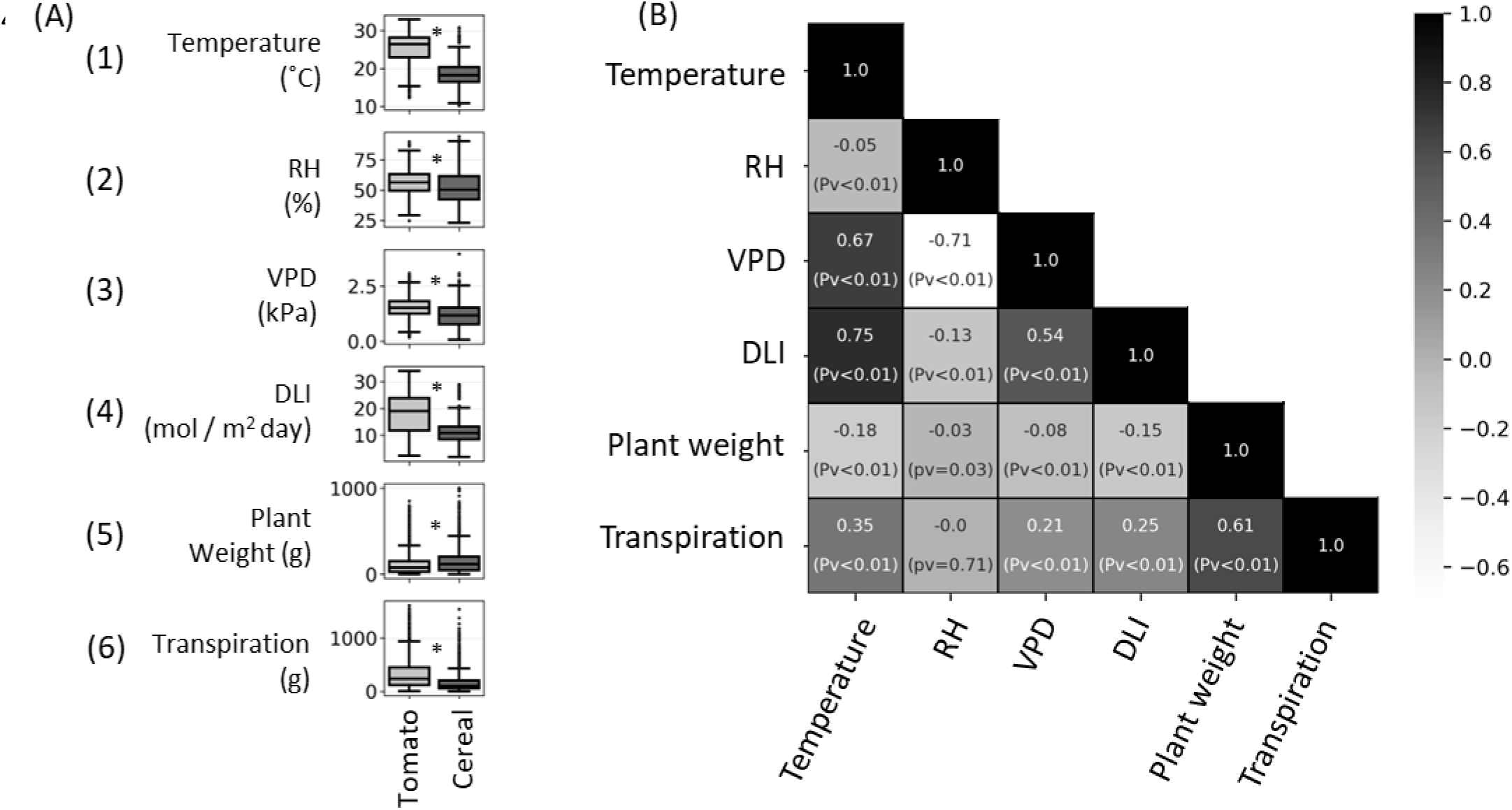
Data visualization. The data was collected between June 2018 and February 2022, by the lysimeter system, and preprocess to daily values. (A) Data variety between tomato and cereals crops (3975 / 2140 observations respectively). Box and Whisker of daily (A1) temperature, (A2) RH, (A3) VPD, (A4) DLI, (A5) plant weight and (A6) plant transpiration (for expansion see Figure S1). Asterisks indicate statistical difference, t test; Pv<=0.001. (B) Pearson correlation coefficient chart of continuous features: Temperature, RH, VPD, DLI, plant weight and transpiration (for expansion see Figure S1). P-values (Pv) are presented in bracket.

Despite cereals having a higher average plant weight (148.67 g) compared to tomatoes (111.54 g), their transpiration rates were significantly lower than those of tomatoes (average 160.24 g/day and 318.32 g/day respectively; Pv <0.001; Fig. 4A5, 4A6). This comparative analysis underscores the distinct environmental adaptabilities and physiological responses of these crops under varying seasonal conditions.

The data, presents a relatively high correlation between DLI and temperature (r = 0.75) and a minor correlation between transpiration and plant weight (r = 0.61; Fig. 4B). VPD is relatively corelated to temperature (r = 0.67), RH (r = −0.71) and DLI (r = 0.54) but not to transpiration (r = 0.21; Fig. 4B).

### 3.1 Model precision and accuracy

Machine learning models were trained using the features: plant weight, temperature, RH, VPD, DLI, plant type, and soil type to predict daily transpiration (Table 1.). Hyperparameters for all models were finely tuned using a Cross-Validation Grid Search on 90% of the training dataset.

Our final models underwent rigorous evaluation through a 10-fold cross-validation process. This involved partitioning the dataset into 10 subsets, with 90% allocated for training and the remaining 10% for testing, repeated across 10 iterations. Such meticulous methodology facilitated comprehensive assessment under diverse training scenarios. To ensure robust performance evaluation, we conducted statistical analyses, including Root Mean Square Error (RMSE) and R² calculations across the 10 folds. Notably, while the Decision Tree model displayed respectable performance, the models Random Forest, XGBoost, and Neural Network showcased significant accuracy (Pv < 0.05, Table 3). Random Forest with the mean R² of 0.88 and an RMSE of 86.93. Similarly, XGBoost and Neural Network models yielded competitive mean R² values of 0.89with RMSE values of 85.19 and 84.63, respectively. Neural Network, however, distinguished itself as the slowest model, completing the 10-fold cross-validation in 9.66 min (Pv < 0.05, Table 3).

### 3.2 Testing the models using hold out experiments

Randomly selected experiments were separated from the rest of the data to serve as a validation dataset for testing the models’ performance on unseen data. The hold out data structure was different from the data used to tune and train the models; with a lower temperature (T test = 14, p ≤ 0.001), different ratio of Tomato: Cereal (Chi-square = 94, Pv < 0.001), and a higher correlation between plant weight and transpiration (Table S1, Fig. S1). Moreover, the distribution of transpiration values in the holdout data significantly differed from those in the training dataset (Pv <0.001). Notably, the upper quartiles of transpiration in the training and holdout datasets were 379g and 255g respectively (Fig. S5 and 5A). When the trained models were applied to this new dataset, the Random Forest model demonstrated high accuracy in predicting transpiration, with a correlation coefficient (r) value of 0.91 (Fig. 5B). The residual plot (Fig. 5C) shows that the model’s residuals are mostly centred around zero, indicating good predictive performance. Note that the distribution is wider where the transpiration is higher. The XGBoost exhibited the highest R^2^ (0.82) and the lowest RMSE (88.41 g) among all models (Table 4, N.S.). The Random Forest Regressor model also performed well (R^2^= 0.81, RMSE = 90.77 g). Conversely, the Neural Network exhibited higher errors (N. S.), with an RMSE of 106 g and an MAE of 79.11 g. On the holdout data, Tukey HSD test revealed no significant differences between models (ANOVA: F= 1.7, Pv = 0.15).

**Figure 5:**
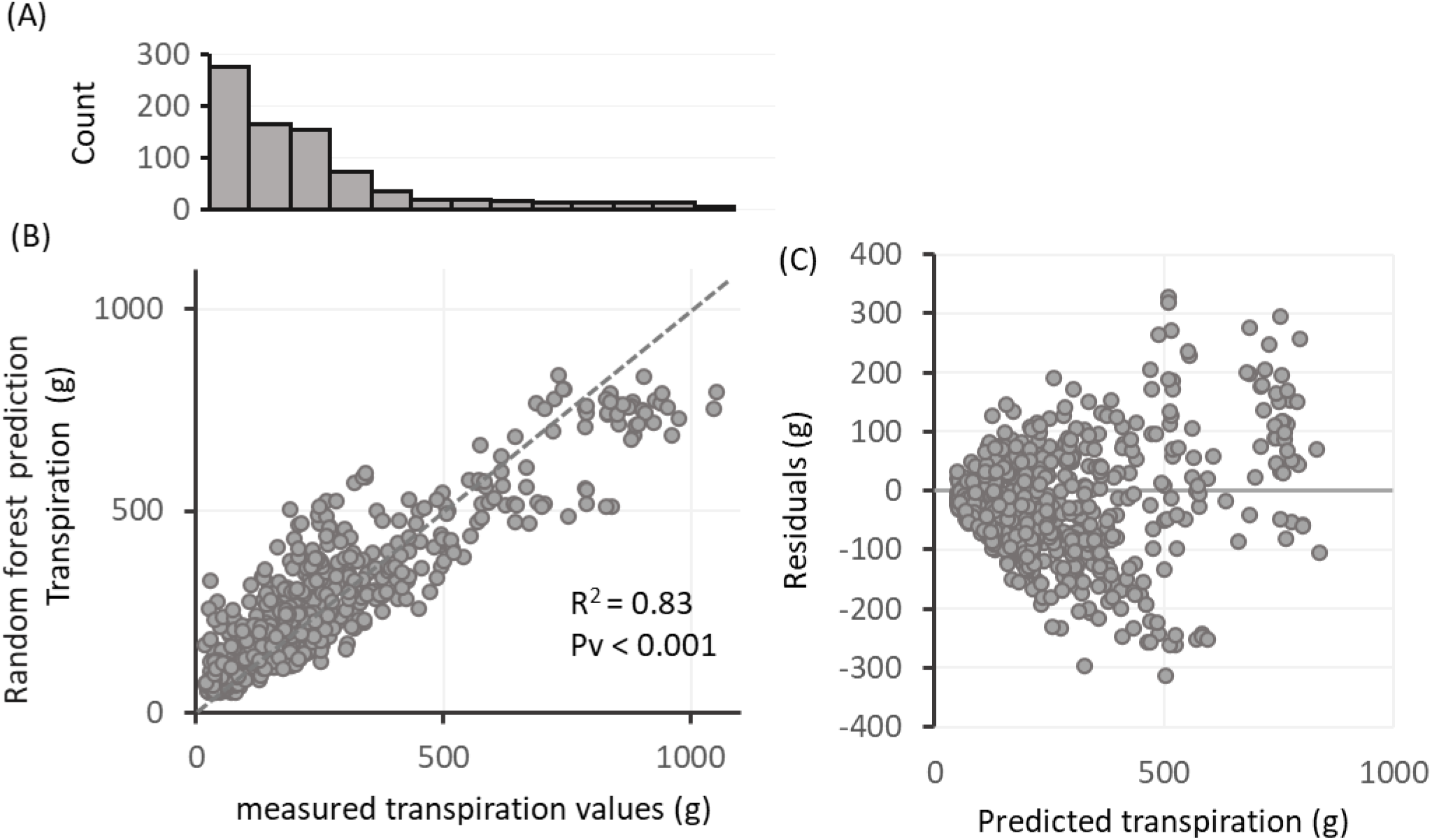
Random forest model evaluation on five holdout experiments (A) Histogram distribution of daily transpiration measured in the holdout data. (B) Goodness of fit for the Random Forest model predictions, comparing predicted against observed values. The dashed gray line represents an exact fit (Y=X), indicating where the model’s predictions perfectly match the observations. The R^2^ and the associated p-value (Pv) are stated on the graph to indicate the strength and significance of the relationship. (C) Residuals (observed minus predicted values) plotted against predicted values to assess any systematic deviations from the model. Higher values, which are less frequent as shown in (A), exhibit increased scatter and reduced correlation, as illustrated in (B) and (C).

Hyperparameter tuning (Table 2) slightly improved the XGBoost model performance on the holdout data, increasing R² from 0.807 to 0.81. To assess whether the tuning prosses improvement was statistically meaningful, we tested the tuning performance across 21 different random seeds, each resulting in a different holdout set. For each subset, we compared model performance with and without hyperparameter tuning. There was no significant difference between the untuned and tuned models (RMSE difference of 0.19 g, paired t test *Pv*= 0.63; Fig. S6).

### 3.2 Model Performance on Independent Greenhouse and Growth Room Experiments

To test the generalizability of our models, we evaluated them on two independent datasets: one from an externally operated greenhouse in Tel Aviv, and one from our controlled growth room (Room 101). The Tel Aviv facility uses the same platform but is operated by a different team and follows independent experimental procedures, while Room 101 provides a distinct indoor environment managed by our own research group. For each experiment, a representative plant was selected to evaluate the models. For more details see Methods Section 2.10.

The models showed good performance on tomato plants across both sites. In Tel Aviv, the Random Forest model achieved an R² of 0.71, while in Room 101 it reached 0.76, with lower RMSE in the latter. For cereal plants in Tel Aviv, XGBoost achieved the lowest RMSE (58.12 g), but the R² was only 0.33, indicating less consistency in capturing variability.

### 3.3 Feature importance

Feature importance was assessed through various techniques. For tree-based models, impurity metrics were employed, permutation and SHAP values were used. Interestingly, plant weight and temperature were identified as crucial predictors for transpiration (Fig. 6A) and the SHAP feature importance test (Fig. 6B) on the Random Forest model predicting transpiration. The horizontal spread of dots for each feature indicates the range of SHAP values, with a wider spread denoting greater impact variability on model output. Color signifies feature value magnitude, with red indicating high and blue indicating low values. Plant weight, temperature, and potting media are the most influential features in predicting transpiration, as evidenced by the wider spread of SHAP values, indicating a stronger effect on model predictions. In contrast, VPD and DLI show a narrow spread with randomly distributed colors, suggesting a small and uniform influence on model output. Divergent color patterns in potting media reveal varying impacts on predictions, with soil (encoded as 1) increasing transpiration and sand (encoded as 0) decreasing it. Furthermore, the cereal plant type (encoded as 1) is associated with reduced transpiration, while the tomato (encoded as 0) contributes to increased transpiration.

**Figure 6:**
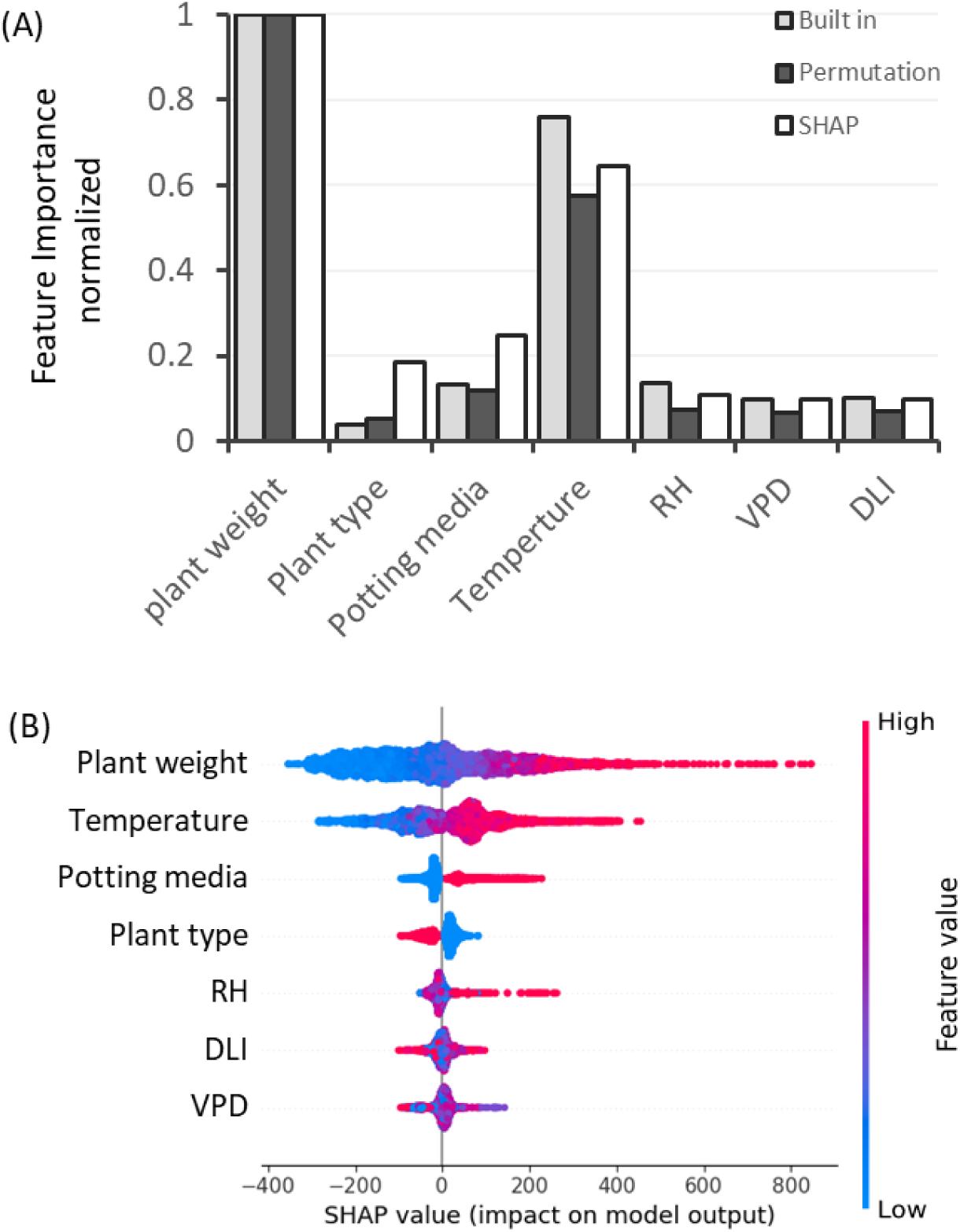
Feature importance contribution to the model prediction accuracy. Plant weight and temperature have a high importance overall. Testing feature significance in **Random Forest** model; (A) Feature importance using built in impurity test (gray), permutation (black) and SHAP values (white). Y axes are the feature importance scores normalized by a Min-Max scaling method, High score is given to the more influential features. (B) SHAP presentation of Feature importance. Every observation has one dot in each row. The position of the dot on the x-axis is the impact of that feature on the model’s prediction for the observation, and the color of the dot represents the value of that feature for the observation. A larger spread of dots suggests that the feature has a more significant impact on the model’s predictions.

### 3.4 Holdout data splitting methods

In this section, we assess the robustness of our data analysis using three distinct partitioning strategies: temporal splits, greenhouse-based splits, and random experiment-based splits. These methods help us evaluate how reliable is our data and how well our models perform across different timeframes, environmental conditions, and experimental setups.

For data partitioning, the temporal split involved segregating data by leaving out a different year for each iteration, using the remaining years for training. This approach allowed us to assess the model’s ability to adapt to temporal variations. The greenhouse-based split used data from one greenhouse for training and testing, while data from another was held out, enabling evaluation of model generalizability across different greenhouse environments. In the random experiment-based split, experiments were randomly selected as holdouts, with the remaining used for training, aiming to gauge the model’s robustness across diverse experimental setups. R^2^ scores analysis indicated no significant performance differences among the splitting methods: temporal split scored approximately 0.54 (median 0.65), random experiment-based split achieved a mean of 0.62(median 0.60), and greenhouse-based split averaged 0.62 (Fig. 7; Welch’s test: F = 0.68, Pv = 0.53). Notably, variations in the random seed introduced considerable variability in the randomly selected holdout sets, while changes across different years also added diversity.

**Figure 7:**
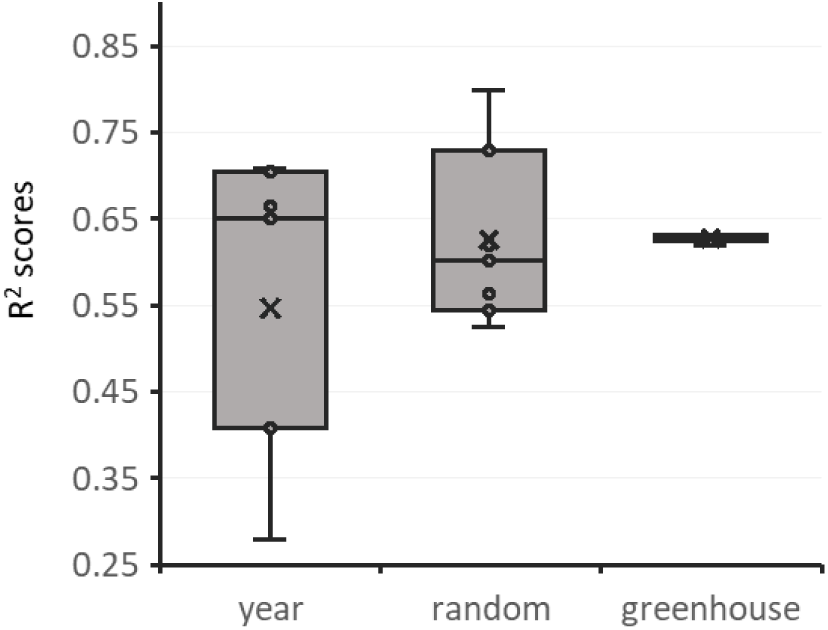
Comparing Data Splitting Methods for Holdout Data Set. The distribution of R2 scores for a tuned random forest model, evaluated using three distinct data splitting methods. The “Year” method separates the data chronologically, training on observations from six years while holdout data was from one excluded year, repeatedly applied across all available years. The “Random” method involves randomly selecting entire experiments as the holdout dataset. Different random seeds lead to varied holdout experiments, resulting in a broad range of scores. The “Greenhouse” method uses data from the main greenhouse for training and data from a secondary, different-sized greenhouse for testing.

## 4. Discussion

In this study, we explored the dynamic of daily transpiration in well-irrigated plants using advance machine learning models. Our unique dataset, collected over seven years, included high-resolution physiological traits and meteorological measurements from two greenhouses. Our findings indicate that our ML models, particularly Random Forest and XGBoost, showed respectable performance in predicting daily transportation. Additionally, both SHAP and feature importance scores revealed that plant weight and daily mean temperature are the most influential factors in these predictions. This discussion addresses these findings and their potential applications in agriculture.

### 4.1. Seasonal Environmental Effects on Plant Weight and Transpiration

Fig. 4 highlights the distinct environmental parameters experienced by summer tomato and winter cereal, as well as the significantly higher daily transpiration of the tomatoes. Interestingly, the plant weight (Fig. 4A5) of cereals was significantly higher than that of tomatoes, with averages of 148.67 g and 111.54 g, respectively. This result was unexpected, given the common correlation between larger annual plant size—which is closely related to transpiring leaf area (Halperin et al., 2017) —and higher transpiration rates (Fig. 4B; Transpiration-Plant Weight; r=0.6). However, this discrepancy highlights how seasonal environmental variations can influence transpiration.

### 4.2. Model performance and Accuracy

The effectiveness of ML models in predicting daily transpiration was tested. The tuned models were trained and evaluated using 10-fold cross-validation. Random Forest and XGBoost performed similarly, with XGBoost being 7 times faster (Table 3). The XGBoost model achieved an RMSE of 85.19 g with and R^2^=0.89. Neural Networks had comparable accuracy but much longer runtimes

Testing our model on a holdout dataset, to further validate our model accuracy and generalizability. The XGBoost model demonstrates the strongest performance in predicting the daily transpiration (R^2^ = 0.82, RMSE = 88.41g). Although there is no significant difference in the residuals between the XGBoost and the Random Forest models.

Interestingly, although hyperparameter tuning (Table 2) improved model performance slightly, testing tuning impact across different random seeds revealed that the improvement was not statistically significant on the holdout data (Fig. S6). This lack of consistent performance gain may be due to the model already being well-calibrated with its default parameters, the limited benefit of tuning given the available feature set, or potential overfitting to the training data during the tuning process.

Our results align with other ML models that have predicted actual transpiration. For instance, Amir et al. (2021) reported that their Random Forest model had R² = 0.81, when predicting sap flow in cherry tomatoes Similarly, Fan et al. (2021) achieved R² values from 0.816 to 0.95, using sap flow data from maize. Ohana-Levi et al. (2020) got R² values of up to 0.90 when predicting grapevine water consumption using drainage lysimeter data. Pagano et al. (2023) present a table summarizing several studies that predicted actual transpiration, with an average maximum R^2^ of 0.87 for these predictions. We propose that the accuracy of our models (R^2^ = 0.81-0.89) can be attributed to the precision of our transpiration measurements, the size of our dataset, and the specific features used during model training (Table. 1). Overall, these findings validate the possibility of using ML models to predict daily transpiration.

Testing our models on other experimental setups, demonstrate the models practical application and external validity in independent environments (Table 5). This external validation provides meaningful insights into the robustness, consistency, and adaptability of our models under varying conditions. Notably, the optimal model differed depending on the plant type and environment. For tomato plants grown in both the Tel Aviv greenhouse and our controlled growth room (Room 101), the Random Forest model outperformed others, achieving R² values of 0.71 and 0.76, with corresponding RMSE values of 81.38 g and 59.58 g, respectively. These results are consistent with the strong performance observed in our holdout experiments (see Fig. 5), reinforcing the model’s reliability. In contrast, for cereal plants grown in Tel Aviv, the XGBoost model demonstrated the lowest RMSE (58.12 g), suggesting high predictive precision. However, the relatively low R^2^ of 0.33 indicated that, while the model’s predictions were close in absolute terms (low RMSE), they did not align well with the variation in actual transpiration values – suggesting that the model failed to explain the full range of responses observed in the cereal plants. This may be due to the lower representation of cereal data (2140 observation versus 3975 for tomato) and the narrower range of environmental condition under which cereal were grown, as evidence in Fig. 4A (temperature, and DLI) which show reduce variability for cereal compared to tomatoes. This divergence in model performance across plant types and conditions emphasizes the importance of model selection tailored to specific experimental contexts

**Table 5:**
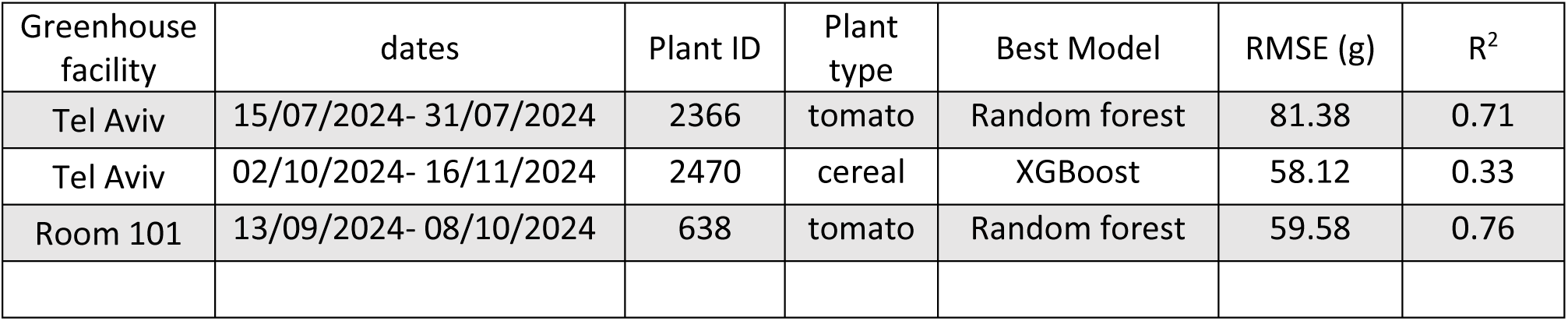
Model Performance on Independent Greenhouse Experiments Using the Model Testing App.

| Greenhouse facility | dates | Plant ID | Plant type | Best Model | RMSE (g) | R<sup>2</sup> |
| --- | --- | --- | --- | --- | --- | --- |
| Tel Aviv | 15/07/2024- 31/07/2024 | 2366 | tomato | Random forest | 81.38 | 0.71 |
| Tel Aviv | 02/10/2024- 16/11/2024 | 2470 | cereal | XGBoost | 58.12 | 0.33 |
| Room 101 | 13/09/2024- 08/10/2024 | 638 | tomato | Random forest | 59.58 | 0.76 |

We suggest that future work should expand the quantity and variability of our labeled datasets by periodically pooling standardized data from additional lysimeter installations. This would enhance model robustness and enable validation across a broader range of crops and climatic conditions. As this global dataset matures, it may become necessary to develop a simple calibration factor— analogous to the FAO-56 crop coefficient—to harmonize measurements across different systems.

### 4.3 Key factors Influencing transpiration

Artificial Intelligence models are often considered “black boxes” due to their complexity, consideration of multiple parameters, and the intricate statistical calculations involved in their algorithms (Azodi et al., 2020). Feature importance analysis enables us to understand the strength of each feature in the model’s predictions and in the transpiration processes.

Our Feature importance analysis indicates that **plant weight** plays the most significant role in predicting daily transpiration (Fig. 6). We find this result very rational as the correlation between plant biomass and transpiration is well known (Lambers et al., 2008; Lazar, 2003). As larger plants have more leaves (and thus more stomata) leading to more water loss. Similarly, other studies have found that canopy size in olives (Sperling et al., 2023) and plant height in maize (Z. Chen et al., 2020) are crucial factors. Additionally, leaf area index has been effectively used in other studies to predict sap flow (Balasubramanian & Thirugnanam, 2023), or grapevine water consumption(Ohana-Levi et al., 2020) highlighting the broader applicability of leaf and plant size in transpiration models.

While direct measurement of plant weight may not be practical in large-scale field applications, it serves as an important physiological proxy for transpiration in controlled settings. Future work could explore remote sensing or image-based methods as scalable alternatives to integrate plant size metrics into transpiration models.

While plant age might replace plant weight for ease, as higher transpiration rates are often observed during late vegetative stages (Grulke & Retzlaff, 2001; Juárez-López et al., 2008). However, in our data, the correlation between plant age and transpiration is weak (days-transpiration; r=0.38, Fig. S1). This due to inconsistencies in documenting plant age in our database, including it as a feature may introduce bias.

**Vapor pressure deficit (VPD)** integrates the effects of both temperature and relative humidity (RH), with the literature often treating them as equally influential on transpiration. However, these two environmental factors not only interact reciprocally—such that a rise in temperature typically coincides with a decrease in RH, and vice versa—but also have distinct direct impacts on plant physiology. For example, while RH primarily affects the atmospheric demand for water vapor, temperature directly influences enzymatic activity and metabolic processes within the plant. As a result, separating the independent contributions of temperature and RH to transpiration responses is challenging. Although VPD has traditionally been considered a key factor influencing stomatal conductance and overall plant transpiration (Song et al., 2022; Tanny, 2013; Zhou et al., 2019) several studies have acknowledged the difficulty of disentangling these interdependent drivers (Bunce, 2006; Grossiord et al., 2020), often focusing on VPD as the primary determinant of water loss rather than parsing the relative roles of its components. Against this backdrop, the findings from this work indicate that temperature exerts a greater influence than RH.

In fact, **temperature** was the second most important factor in our feature importance hierarchy, making it our most critical environmental predictor of daily transpiration (Fig. 6). This aligns with its previously observed effects on physiological processes, as high temperatures drive transpiration by increasing water vapor pressure (Aschale et al., 2023; W. Liu et al., 2020). Additionally, it directly affects the physiological responses of plants, such as stomatal conductance (Haijun et al., 2015). Some researchers even found temperature to be the most critical factor influencing evapotranspiration (ET0), especially in spring and summer season when it contributes to as much as 46% of the variance of ET0 (Aschale et al., 2023; W. Liu et al., 2020). Nevertheless, our results suggest that temperature alone may have a much stronger effect than RH. Thus, the high feature importance assigned to temperature in our model may indicate its predominant role in controlling transpiration, independent of the relative humidity component of VPD. Further studies are needed to confirm and clarify this distinction.

**Potting media** is the third most important feature (Fig. 6B). SHAP analysis showed that potting media are completely separated, and sand (blue) has negative effect on the transpiration compared to the soil (red). This is due to differences in water availability, as soil typically retains more water than sand, leading to higher transpiration in plants grown in soil(Cai et al., 2024).

We were surprised to find that **plant type** had a relatively low importance in predicting transpiration (Fig. 6A). Genotype by Environment (GxE) interactions suggest that different crops like tomatoes and cereals, with potential variations in stomatal density, would influence transpiration rates (Des Marais et al., 2013; Fournier-Level et al., 2011). A possible explanation is that the model may identify plant type indirectly through weight (Fig. 4A5, Fig. 6A). However, if the model’s assessment holds, it could signal a ground-breaking insight into the plant transpiration prediction.

The low importance score of the **Daily Light Integral (DLI)** was unexpected, as it contradicts the common understanding of the high significance of solar radiation (Tanny, 2013). It is possible that in indirect models based on eddy covariance (Pagano et al., 2023) and FAO56 estimation (Başağaoğlu et al., 2021), radiation plays a more crucial role, whereas in models using direct measurements such as sap and lysimeters (current article), its importance may be less pronounced. Another consideration is that daily averaging might diminish the perceived importance of light, while more granular data collected at hourly or minute intervals may reveal its greater significance (as in these works: Kiraga et al., 2023; Li et al., 2020).

Future studies should explore the use of more granular data, such as hourly or minute-level measurements, instead of daily averages, to better capture the influence of all meteorological parameters on transpiration. Daily averaging can obscure the dynamic effects of factors like solar radiation, temperature, and humidity, potentially underestimating their true importance. By analyzing shorter time intervals, researchers may uncover stronger correlations between these variables and transpiration, providing a clearer understanding of their interactions.

It is important to note that soil water content, ambient CO₂ concentration, and wind speed were maintained at non-stress or quasi-steady levels during the experiments (see Materials and Methods) and were therefore not included as predictors. Because these variables are known to influence transpiration, future work should broaden the set of independent drivers and incorporate their dynamic behaviour into the machine-learning framework and model training.

We suggest that future elaboration and integration of additional ambient variables (such as CO₂ concentration, wind speed, soil electrical conductivity (EC), and soil water content) should be incorporated into the methodological framework proposed here. These factors exert direct, indirect, and interactive effects on plant behavior, and their inclusion could improve both the resolution of hierarchical environmental controls on whole-plant transpiration and the accuracy of transpiration prediction across a wider range of environmental conditions and growth systems.

### 4.4 Splitting methods

Splitting the data into train and validation sets can significantly impact model performance and generalizability (Shi et al., 2022). We explored three distinct data sampling methods: year (temporal), random, and greenhouse (spatial). Although the models’ accuracy results didn’t show a significant difference between them, each method offered unique insights into the data strengths and the models’ ability to accurately predict transpiration (Fig. 7). The year split had a median accuracy (R^2^) of 0.65, impacted by greenhouse differences, equipment aging, and yearly meteorological variation. Random sampling yielded a median accuracy of 0.60, with a broader variation. Greenhouse-based splitting achieved 0.63, as the model effectively predicted transpiration in the secondary greenhouse using data from the main one, suggesting good generalizability across similar coupled setups.

### 4.5. Future Applications and Model-Based Decision Support

In this study, we demonstrated that machine learning models can achieve respectable predictive performance using only a small number of input features. Moreover, the feature importance analysis reveals that the model’s internal weighting aligns well with established physiological drivers of transpiration, reinforcing both its predictive value and biological relevance.

Future research should prioritize expanding the diversity, resolution, and temporal coverage of environmental and physiological data by integrating advanced sensors for continuous spatiotemporal monitoring. This will enhance our ability to model dynamic plant-environment interactions with greater accuracy. In addition to improving prediction, AI models may enable early detection of suboptimal plant behavior and stress responses, supporting proactive decision-making in precision agriculture. These future models are particularly valuable for disentangling multifactorial influences, such as the overlapping yet distinct effects of temperature and humidity, and for revealing non-intuitive interactions that might be overlooked in traditional analyses. Identifying the most influential variables also enables the strategic use of low-cost sensors in data-driven irrigation and environmental control systems. As datasets expand and model accuracy improves, predictions may increasingly rely on ambient measurements alone, with lysimeters serving primarily as initial calibration references. Ultimately, physiology-informed ML models can support scalable, sensor-efficient, and predictive crop management, bridging fundamental understanding with real-world implementation.

## Acknowledgments

This research was partly supported by the Shoenberg Research Center for Agricultural Science (Grant #3175006230), the Israel Science Foundation (Grant No. 1043/20), and the Israeli Committee for Budgeting for their support in the research of sustainable agriculture and food security at the Israeli Center for Digital Agriculture. We thank Dr. Nir Sade (Tel Aviv University) for generously sharing data from his PlantArray phenotyping system.

## Competing interests

None declared.

## Author contributions

SF and MM designed the study. SF collected the data and performed the data analyses. TN conducted the initial coding and data cleaning. SF and NA adjusted the workflow and tested the statistical models. SF wrote the first draft of the manuscript. MM (Corresponding Author) oversaw manuscript revisions, project management, and secured funding. SF, NA and MM contributed substantially to revisions.

## Data availability

Data available on request due to privacy/ethical restrictions

## Supporting Information

**Supplementary Methods S1:** Description of the systematic literature search and filtering procedure for identifying machine learning studies focused on transpiration and evapotranspiration modeling.

**Supplementary Table 1:** Description of the training and holdout datasets used for model training and evaluation.

**Supplementary Figure 1:** Correlation charts of the training and holdout datasets, showing the Pearson correlation coefficients.

**Supplementary Figure 2:** Environmental conditions and experimental setup in the indoor growth room used for external model validation.

**Supplementary Figure 3:** Distributions of environmental and physiological parameters by soil type and crop type.

**Supplementary Figure 4:** SHAP (SHapley Additive exPlanations) analysis showing the impact of features on transpiration predictions.

**Supplementary Figure 5:** Box and whisker plots illustrating the distribution of transpiration in the training and holdout datasets.

**Supplementary Figure 6:** Evaluation of model performance with and without hyperparameter tuning using random forest models. The analysis shows no statistically significant improvement in RMSE after tuning.

## Notes

### Competing Interest Statement

The authors have declared no competing interest.

### Summary of Updates

We expanded the literature review to include a systematic analysis of recent studies utilizing machine learning for transpiration prediction. Several new figures were added to illustrate the experimental setup, greenhouses, and data architecture. External validation results were incorporated using independently collected datasets from a separate greenhouse and a controlled growth room environment, demonstrating the robustness and generalizability of the model. The greenhouse feature was removed, and the dataset properties, results, and hyperparameter tuning procedures were updated accordingly. Minor text clarifications and structural edits were made throughout to improve readability and coherence.

## References

Abadi, M., Agarwal, A., Barham, P., Brevdo, E., Chen, Z., Citro, C., Corrado, G. S., Davis, A., Dean, J., Devin, M., Ghemawat, S., Goodfellow, I., Harp, A., Irving, G., Isard, M., Jia, Y., Jozefowicz, R., Kaiser, L., Kudlur, M., … Research, G. (2016). TensorFlow: Large-Scale Machine Learning on Heterogeneous Distributed Systems. ArXiv Preprint. www.tensorflow.org.

Amani, S., & Shafizadeh-Moghadam, H. (2023). A review of machine learning models and influential factors for estimating evapotranspiration using remote sensing and ground-based data. Agricultural Water Management, 284, 108324. 10.1016/J.AGWAT.2023.108324

Amir, A., Butt, M., & Van Kooten, O. (2021). Using Machine Learning Algorithms to Forecast the Sap Flow of Cherry Tomatoes in a Greenhouse. IEEE Access, 9, 154183–154193. 10.1109/ACCESS.2021.3127453

Anapalli, S. S., Ahuja, L. R., Gowda, P. H., Ma, L., Marek, G., Evett, S. R., & Howell, T. A. (2016). Simulation of crop evapotranspiration and crop coefficients with data in weighing lysimeters. Agricultural Water Management, 177, 274–283. 10.1016/J.AGWAT.2016.08.009

Aschale, T. M., Peres, D. J., Gullotta, A., Sciuto, G., & Cancelliere, A. (2023). Trend Analysis and Identification of the Meteorological Factors Influencing Reference Evapotranspiration. Water 2023, Vol. 15, Page 470, 15(3), 470. 10.3390/W15030470

Averbuch, N., & Moshelion, M. (2024). Evaluating Evapotranspiration in a Commercial Greenhouse: A Comparative Study of Microclimatic Factors and Machine-Learning Algorithms. BioRxiv, 2024.01.11.575151. 10.1101/2024.01.11.575151

Azodi, C. B., Tang, J., & Shiu, S. H. (2020). Opening the Black Box: Interpretable Machine Learning for Geneticists. Trends in Genetics, 36(6), 442–455. 10.1016/J.TIG.2020.03.005/ASSET/2F2EDEAA-DCFE-4EB8-A287-6955F2A064F0/MAIN.ASSETS/GR3.JPG

Balasubramanian, H. K., & Thirugnanam, H. (2023). Neural Networking to Predict Sap Flow Using AI-Synthesized Relative Meteorological Data. 2023 3rd International Conference on Intelligent Technologies, CONIT 2023. 10.1109/CONIT59222.2023.10205886

Ball, J. T., Woodrow, I. E., & Berry, J. A. (1987). A model predicting stomatal conductance and its contribution to the control of photosynthesis under different environmental conditions. Progress in Photosynthesis Research, 90(1), 221–224. 10.1007/978-94-017-0519-6_48

Başağaoğlu, H., Chakraborty, D., & Winterle, J. (2021). Reliable Evapotranspiration Predictions with a Probabilistic Machine Learning Framework. Water 2021, Vol. 13, Page 557, 13(4), 557. 10.3390/W13040557

Ben-Asher, J., Garcia Y Garcia, A., & Hoogenboom, G. (2008). Effect of high temperature on photosynthesis and transpiration of sweet corn (Zea mays L. var. rugosa). Photosynthetica, 46(4), 595–603. 10.1007/S11099-008-0100-2

Breiman, L. (2001). Random forests. Machine Learning, 45(1), 5–32. 10.1023/A:1010933404324/METRICS

Breiman, L., Friedman, J. H., Olshen, R. A., & Stone, C. J. (1984). Classification and regression trees. In Classification and Regression Trees (1st ed.). Routledge. 10.1201/9781315139470

Bunce, J. A. (2006). How do leaf hydraulics limit stomatal conductance at high water vapour pressure deficits? Plant, Cell & Environment, 29(8), 1644–1650. 10.1111/J.1365-3040.2006.01541.X

Cai, G., König, M., Carminati, A., Abdalla, M., Javaux, M., Wankmüller, F., & Ahmed, M. A. (2024). Transpiration response to soil drying and vapor pressure deficit is soil texture specific. Plant and Soil, 500(1–2), 129–145. 10.1007/s11104-022-05818-2

Chen, T., & Guestrin, C. (2016). XGBoost: A Scalable Tree Boosting System. Proceedings of the 22nd ACM SIGKDD International Conference on Knowledge Discovery and Data Mining, 785–794. 10.1145/2939672.2939785

Chen, Z., Sun, S., Wang, Y., Wang, Q., & Zhang, X. (2020). Temporal convolution-network-based models for modeling maize evapotranspiration under mulched drip irrigation. Computers and Electronics in Agriculture, 169, 105206. 10.1016/J.COMPAG.2019.105206

Chollet, F. (2015). Keras: Deep Learning for humans. https://keras.io/

Cutler, A., & Zhao, G. (2001). PERT-perfect random tree ensembles. https://www.researchgate.net/publication/268424569

Dalal, A., Shenhar, I., Bourstein, R., Mayo, A., Grunwald, Y., Averbuch, N., Attia, Z., Wallach, R., & Moshelion, M. (2020). A telemetric, gravimetric platform for real-time physiological phenotyping of plant–environment interactions. Journal of Visualized Experiments, 162(e61280), 1–28. 10.3791/61280

de Meneses, K. C., Aparecido, L. E. D. O., de Meneses, K. C., & de Farias, M. F. (2020). Estimating Potential Evapotranspiration in Maranhão State Using Artificial Neural Networks. Revista Brasileira de Meteorologia, 35(4), 675–682. 10.1590/0102-77863540072

Des Marais, D. L., Hernandez, K. M., & Juenger, T. E. (2013). Genotype-by-Environment Interaction and Plasticity: Exploring Genomic Responses of Plants to the Abiotic Environment. Annual Review of Ecology, Evolution, and Systematics, 44(1), 5–29. 10.1146/annurev-ecolsys-110512-135806

Dixon, M., & Grace, J. (1984). Effect of wind on the transpiration of young trees. Annals of Botany, 53(6), 811–819. 10.1093/OXFORDJOURNALS.AOB.A086751

Fan, J., Zheng, J., Wu, L., & Zhang, F. (2021). Estimation of daily maize transpiration using support vector machines, extreme gradient boosting, artificial and deep neural networks models. Agricultural Water Management, 245, 106547. 10.1016/J.AGWAT.2020.106547

Ferreira, L. B., da Cunha, F. F., de Oliveira, R. A., & Fernandes Filho, E. I. (2019). Estimation of reference evapotranspiration in Brazil with limited meteorological data using ANN and SVM – A new approach. Journal of Hydrology, 572, 556–570. 10.1016/J.JHYDROL.2019.03.028

Fournier-Level, A., Korte, A., Cooper, M. D., Nordborg, M., Schmitt, J., & Wilczek, A. M. (2011). A Map of Local Adaptation in Arabidopsis thaliana. Science, 334(6052), 86–89. 10.1126/science.1209271

Fukushima. (1969). Visual Feature Extraction by a Multilayered Network of Analog Threshold Elements. IEEE Transactions on Systems Science and Cybernetics, 5(4), 322–333. 10.1109/TSSC.1969.300225

Fürnkranz, J. (2011). Decision Tree. Encyclopedia of Machine Learning, 263–267. 10.1007/978-0-387-30164-8_204

Geller, G. N., & Smith, W. K. (1982). Influence of leaf size, orientation, and arrangement on temperature and transpiration in three high-elevation, large-leafed herbs. Oecologia, 53(2), 227–234. 10.1007/BF00545668/METRICS

Gosa, S. C., Lupo, Y., & Moshelion, M. (2019). Quantitative and comparative analysis of whole-plant performance for functional physiological traits phenotyping: New tools to support pre-breeding and plant stress physiology studies. Plant Science, 282, 49–59. 10.1016/J.PLANTSCI.2018.05.008

Grossiord, C., Buckley, T. N., Cernusak, L. A., Novick, K. A., Poulter, B., Siegwolf, R. T. W., Sperry, J. S., & McDowell, N. G. (2020). Plant responses to rising vapor pressure deficit. New Phytologist, 226(6), 1550–1566. 10.1111/NPH.16485;PAGE:STRING:ARTICLE/CHAPTER

Grulke, N. E., & Retzlaff, W. A. (2001). Changes in physiological attributes of ponderosa pine from seedling to mature tree. Tree Physiology, 21(5), 275–286. 10.1093/TREEPHYS/21.5.275

Haijun, L., Cohen, S., Lemcoff, J. H., Israeli, Y., & Tanny, J. (2015). Sap flow, canopy conductance and microclimate in a banana screenhouse. Agricultural and Forest Meteorology, 201, 165–175. 10.1016/j.agrformet.2014.11.009

Halperin, O., Gebremedhin, A., Wallach, R., & Moshelion, M. (2017). High-throughput physiological phenotyping and screening system for the characterization of plant–environment interactions. Plant Journal, 89(4), 839–850. 10.1111/tpj.13425

Harris, C. R., Millman, K. J., van der Walt, S. J., Gommers, R., Virtanen, P., Cournapeau, D., Wieser, E., Taylor, J., Berg, S., Smith, N. J., Kern, R., Picus, M., Hoyer, S., van Kerkwijk, M. H., Brett, M., Haldane, A., del Río, J. F., Wiebe, M., Peterson, P., … Oliphant, T. E. (2020). Array programming with NumPy. Nature 2020 585:7825, 585(7825), 357–362. 10.1038/s41586-020-2649-2

Haykin, S. (2009). Neural networks and learning machines (3rd ed.). Pearson Education India.

Ho, T. K. (1995). Random decision forests. Proceedings of the International Conference on Document Analysis and Recognition, ICDAR, 1, 278–282. 10.1109/ICDAR.1995.598994

Imai, K., & Murata, Y. (1976). Effect of carbon dioxide concentration on growth and dry matter production of crop plants. Effects on leaf area, dry matter, tillering, dry matter distribution ratio and transpiration. . Japanese Journal of Crop Science, 45(4), 598–606.

Iqbal, A., Fahad, S., Iqbal, M., Alamzeb, M., Ahmad, A., Anwar, S., Khan, A., Arif, M., Inamullah, S., Saeed, M., & Song, M. (2020). Hasanuzzaman, M., Tanveer, M. (eds) Salt and Drought Stress Tolerance in Plants. Signaling and Communication in Plants. Springer, Cham. 10.1007/978-3-030-40277-8_4

Juárez-López, F. J., Escudero, A., & Mediavilla, S. (2008). Ontogenetic changes in stomatal and biochemical limitations to photosynthesis of two co-occurring Mediterranean oaks differing in leaf life span. Tree Physiology, 28(3), 367–374. 10.1093/TREEPHYS/28.3.367

Kiraga, S., Peters, R. T., Molaei, B., Evett, S. R., & Marek, G. (2023). Reference Evapotranspiration Estimation Using Genetic Algorithm-Optimized Machine Learning Models and Standardized Penman-Monteith Equation in a Highly Advective Environment. 10.3390/w16010012

Lambers, H., Raven, J. A., Shaver, G. R., & Smith, S. E. (2008). Plant nutrient-acquisition strategies change with soil age. Trends in Ecology and Evolution, 23(2), 95–103. 10.1016/J.TREE.2007.10.008/ASSET/CBF09F7C-765E-4123-ACAB-F9E82000E550/MAIN.ASSETS/GR3.JPG

Landeras, G., Ortiz-Barredo, A., & López, J. J. (2008). Comparison of artificial neural network models and empirical and semi-empirical equations for daily reference evapotranspiration estimation in the Basque Country (Northern Spain). Agricultural Water Management, 95(5), 553–565. 10.1016/J.AGWAT.2007.12.011

Lange, O. L., Lösch, R., Schulze, E. D., & Kappen, L. (1971). Responses of stomata to changes in humidity. Planta, 100(1), 76–86. 10.1007/BF00386887

Lazar, T. (2003). Taiz, L. and Zeiger, E. Plant physiology. 3rd edn. Annals of Botany, 91(6), 750–751. 10.1093/AOB/MCG079

Leibniz, G. W. F. von. (1920). The Early Mathematical Manuscripts of Leibniz (C. I. Gerhardt, Ed.). Courier Corporation. https://books.google.co.il/books?hl=en&lr=&id=tCmp_c3Q9S8C&oi=fnd&pg=PP1&dq=Leibniz,+Gottfried+Wilhelm+Freiherr+von+(1920).%C2%A0The+Early+Mathematical+Manuscripts+of+Leibniz:+Translated+from+the+Latin+Texts+Published+by+Carl+Immanuel+Gerhardt+with+Critical+and+Historical+Notes+(Leibniz+published+the+chain+rule+in+a+1676+memoir).+Open+court+publishing+Company.%C2%A0ISBN%C2%A09780598818461.&ots=eD4SCW8omT&sig=dnpMq4OoEyLVJZDfHxQcw9oL_3U&redir_esc=y#v=onepage&q&f=false

Li, L., Chen, S., Yang, C., Meng, F., & Sigrimis, N. (2020). Prediction of plant transpiration from environmental parameters and relative leaf area index using the random forest regression algorithm. Journal of Cleaner Production, 261, 121136. 10.1016/J.JCLEPRO.2020.121136

Liu, W., Zhang, B., & Han, S. (2020). Quantitative Analysis of the Impact of Meteorological Factors on Reference Evapotranspiration Changes in Beijing, 1958–2017. Water 2020, Vol. 12, Page 2263, 12(8), 2263. 10.3390/W12082263

Liu, X., Xu, C., Zhong, X., Li, Y., Yuan, X., & Cao, J. (2017). Comparison of 16 models for reference crop evapotranspiration against weighing lysimeter measurement. Agricultural Water Management, 184, 145–155. 10.1016/J.AGWAT.2017.01.017

López-Urrea, R., Martín de Santa Olalla, F., Fabeiro, C., & Moratalla, A. (2006). Testing evapotranspiration equations using lysimeter observations in a semiarid climate. Agricultural Water Management, 85(1–2), 15–26. 10.1016/J.AGWAT.2006.03.014

Lundberg, S. M., Allen, P. G., & Lee, S.-I. (2017). A Unified Approach to Interpreting Model Predictions. https://github.com/slundberg/shap

Madhu, M., & Hatfield, J. L. (2014). Interaction of carbon dioxide enrichment and soil moisture on photosynthesis, transpiration, and water use efficiency of soybean. Agricultural Sciences, 5(5), 410–429. 10.4236/AS.2014.55043

Mckinney, W. (2010). Data Structures for Statistical Computing in Python.

Merrick, L., & Taly, A. (2020). The Explanation Game: Explaining Machine Learning Models Using Shapley Values. In A. Holzinger, peter Kieseberg, a min Tjoa, & E. weippl (Eds.), Machine Learning and Knowledge Extraction (pp. 17–38). 10.1007/978-3-030-57321-8_2

Mingers, J. (1989). An empirical comparison of selection measures for decision-tree induction. Machine Learning 1989 3:4, 3(4), 319–342. 10.1007/BF00116837

Molnar, C. (2022). Chapter 9 Local Model-Agnostic Methods | Interpretable Machine Learning. https://christophm.github.io/interpretable-ml-book/local-methods.html

Monteith, J. L. (1965). Evaporation and environment. Symposia of the Society for Experimental Biology, 19, 205–234.

Ohana-Levi, N., Munitz, S., Ben-Gal, A., Schwartz, A., Peeters, A., & Netzer, Y. (2020). Multiseasonal grapevine water consumption – Drivers and forecasting. Agricultural and Forest Meteorology, 280, 107796. 10.1016/J.AGRFORMET.2019.107796

Pagano, A., Amato, F., Ippolito, M., De Caro, D., Croce, D., Motisi, A., Provenzano, G., & Tinnirello, I. (2023). Machine learning models to predict daily actual evapotranspiration of citrus orchards under regulated deficit irrigation. Ecological Informatics, 76, 102133. 10.1016/J.ECOINF.2023.102133

Pedregosa, F., Michel, V., Grisel, O., Blondel, M., Prettenhofer, P., Weiss, R., Vanderplas, J., Cournapeau, D., Pedregosa, F., Varoquaux, G., Gramfort, A., Thirion, B., Grisel, O., Dubourg, V., Passos, A., Brucher, M., Perrot and Édouard and, M., Duchesnay, and Édouard, & Duchesnay, Fré. (2011). Scikit-learn: Machine Learning in Python. Journal of Machine Learning Research, 12(85), 2825–2830. http://jmlr.org/papers/v12/pedregosa11a.html

Penman, H. Latimer. (1948). Natural evaporation from open water, bare soil and grass. Proceedings of the Royal Society, Series A, 193(1032), 120–145.

Pieruschka, R., Huber, G., & Berry, J. A. (2010). Control of transpiration by radiation. Proceedings of the National Academy of Sciences of the United States of America, 107(30), 13372–13377. 10.1073/pnas.0913177107

Quinlan, J. R. (1993). C4.5: Programs for Machine Learning - J. Ross Quinlan - Google Books. Morgan Kaufmann Publishers. https://books.google.co.il/books?hl=en&lr=&id=b3ujBQAAQBAJ&oi=fnd&pg=PP1&dq=Quinlan,+J.+R.+(1993).%C2%A0C4.5:+Programs+for+machine+learning.+San+Mateo:+Morgan+Kaufmann.+&ots=sS1tRPJmC6&sig=d1hJNAHV650dcwNzQbol63WLRno&redir_esc=y#v=onepage&q=Quinlan%2C%20J.%20R.%20(1993).%C2%A0C4.5%3A%20Programs%20for%20machine%20learning.%20San%20Mateo%3A%20Morgan%20Kaufmann.&f=false

Rawson, H. M., Begg, J. E., & Woodward, R. G. (1977). The effect of atmospheric humidity on photosynthesis, transpiration and water use efficiency of leaves of several plant species. Planta, 134(1), 5–10. 10.1007/BF00390086/METRICS

Rosenblatt, F. (1958). The perceptron: A probabilistic model for information storage and organization in the brain. Psychological Review, 65(6), 386–408. 10.1037/H0042519

Shapley, L. S. (1953). 17. A Value for n-Person Games. In H. W. Kuhn & A. W. Tucker (Eds.), Contributions to the Theory of Games (AM-28) (Vol. 2, pp. 307–318). Princeton University Press. 10.1515/9781400881970-018

Shi, H., Luo, G., Hellwich, O., Xie, M., Zhang, C., Zhang, Y., Wang, Y., Yuan, X., Ma, X., Zhang, W., Kurban, A., De Maeyer, P., & Van De Voorde, T. (2022). Evaluation of water flux predictive models developed using eddy-covariance observations and machine learning: a meta-analysis. Hydrology and Earth System Sciences, 26(18), 4603–4618. 10.5194/HESS-26-4603-2022

Song, Y., Jiao, W., Wang, J., & Wang, L. (2022). Increased Global Vegetation Productivity Despite Rising Atmospheric Dryness Over the Last Two Decades. Earth’s Future, 10(7), e2021EF002634. 10.1029/2021EF002634;WGROUP:STRING:PUBLICATION

Sperling, O., Yermiyahu, U., & Hochberg, · Uri. (2023). Linking almond trees’ transpiration to irrigation’s mineral composition by physiological indices and machine learning. Irrigation Science, 41, 487–499. 10.1007/s00271-022-00803-0

Tanny, J. (2013). Microclimate and evapotranspiration of crops covered by agricultural screens: A review. In Biosystems Engineering (Vol. 114, Issue 1, pp. 26–43). 10.1016/j.biosystemseng.2012.10.008

The pandas development team. (2020). pandas-dev/pandas: Pandas. Zenodo. 10.5281/ZENODO.10537285

Virtanen, P., Gommers, R., Oliphant, T. E., Haberland, M., Reddy, T., Cournapeau, D., Burovski, E., Peterson, P., Weckesser, W., Bright, J., van der Walt, S. J., Brett, M., Wilson, J., Millman, K. J., Mayorov, N., Nelson, A. R. J., Jones, E., Kern, R., Larson, E., … Vázquez-Baeza, Y. (2020). SciPy 1.0: fundamental algorithms for scientific computing in Python. Nature Methods 2020 17:3, 17(3), 261–272. 10.1038/s41592-019-0686-2

Xing, L., Cui, N., Liu, C., Zhao, L., Guo, L., Du, T., Zhan, C., Wu, Z., Wen, S., & Jiang, S. (2022). Estimation of daily apple tree transpiration in the Loess Plateau region of China using deep learning models. Agricultural Water Management, 273, 107889. 10.1016/J.AGWAT.2022.107889

Zhou, S., Zhang, Y., Williams, A. P., & Gentine, P. (2019). Projected increases in intensity, frequency, and terrestrial carbon costs of compound drought and aridity events. Science Advances, 5(1). 10.1126/SCIADV.AAU5740/SUPPL_FILE/AAU5740_SM.PDF

