## Supporting Information for "Integrating Load-Cell Lysimetry and Machine Learning for Prediction of Daily Plant Transpiration"

### Supplementary

**Supplementary Methods S1:** Literature Search and Filtering Procedure

To expand our review of machine learning applications in transpiration and evapotranspiration modeling, we conducted a systematic literature search using the Scopus database [https://www.scopus.com/search/form.uri?display=basic#basic]. Our search targeted peer-reviewed journal articles published between 2010 and 2024, limited to the subject areas of Environmental Science, Earth and Planetary Sciences, and Agricultural and Biological Sciences, and filtered by article document type to ensure relevance and scholarly quality.

We used the following Boolean search string, applied to the title, abstract, and keywords fields: ("transpiration" OR "evapotranspiration" OR "ET") AND ("machine learning" OR "artificial intelligence"). The search was restricted to peer-reviewed journal articles published between 2010 and 2024 within the subject areas of Environmental Science, Earth and Planetary Sciences, and Agricultural and Biological Sciences. We limited the document type to articles only and exported the citation information along with abstracts and keywords. The search was conducted on February 25, 2025, and returned 2,061 articles.

To refine the dataset, we implemented a keyword-based filtering pipeline in Python. Abstracts were first preprocessed by converting all text to lowercase and removing entries with missing content. Inclusion criteria required that abstracts mention at least one ML-related term (e.g., machine learning, neural network, random forest, SVM, deep learning), a transpiration-related term (e.g., transpiration, evapotranspiration), and at least one predictive keyword (e.g., predict, forecast, modeling). Articles were excluded if they contained irrelevant terms indicating a focus outside the scope of plant-environment modeling—such as groundwater flow, aquifer modeling, pollution, fire dynamics, yield forecasting, or purely hydrological or geospatial applications. This process reduced the initial set of 2,061 articles to a curated collection of 542 studies that directly addressed machine learning-based prediction of transpiration or evapotranspiration.

Each abstract was further analyzed to identify the type of transpiration measurement methods and environmental features used. We categorized measurement approaches into direct methods (e.g., lysimeters, sap flow sensors, porometry, gas exchange systems) and indirect methods (e.g., remote sensing, eddy covariance, and empirical models like FAO Penman-Monteith). Input features were classified into four main groups: (1) climate and meteorological parameters (e.g., temperature, humidity, solar radiation, wind speed, VPD), (2) vegetation indices (e.g., NDVI, LAI, chlorophyll index), (3) soil characteristics (e.g., soil moisture, water potential, texture), and (4) plant physiological traits (e.g., stomatal conductance, transpiration rate, photosynthesis, growth metrics). The annotated dataset includes information on the type of data used, modeling targets, and methodological approaches, thereby offering a clear overview of the state of the field and key knowledge gaps—particularly the limited use of direct physiological measurements in predictive modeling frameworks.

**Supplementary Table 1:** Dataset description: (A) Training dataset description, (B) Holdout dataset description; the holdout dataset consists of 4 randomly chosen independent experiments.

(A) Training dataset

| | VPD | Temp | RH | DLI | Transpiration | encoded_plant | encoded_soil | plant_weight_process |
|---|---|---|---|---|---|---|---|---|
| count | 5310 | 5310 | 5310 | 5310 | 5310 | 5310 | 5310 | 5310 |
| mean | 1.453118 | 23.28861 | 54.65711 | 16.44557 | 270.0401 | 0.326742 | 0.302637 | 121.0753 |
| std | 0.526248 | 4.979421 | 11.91229 | 7.224102 | 255.0648 | 0.469066 | 0.459443 | 129.9826 |
| min | 0.070639 | 10.20526 | 23.03113 | 1.89396 | 1.45 | 0 | 0 | 1 |
| 25% | 1.124764 | 19.10945 | 46.47273 | 10.68273 | 82.2025 | 0 | 0 | 29.94439 |
| 50% | 1.428718 | 24.24519 | 54.27059 | 15.34428 | 185.835 | 0 | 0 | 82.35008 |
| 75% | 1.79233 | 27.61404 | 62.5496 | 22.47608 | 379.3625 | 1 | 1 | 162.1425 |
| max | 3.963345 | 32.93702 | 94.38441 | 33.98652 | 1617.49 | 1 | 1 | 1003.854 |

(B) Holdout dataset

| | vpd | Temp | RH | DLI | Transpiration | encoded_plant | encoded_soil | plant_weight_process |
|---|---|---|---|---|---|---|---|---|
| count | 805 | 805 | 805 | 805 | 805 | 805 | 805 | 805 |
| mean | 1.215469 | 20.86302 | 56.24715 | 11.94429 | 216.5662 | 0.503106 | 0 | 147.3449 |
| std | 0.44616 | 4.307387 | 12.88273 | 5.443994 | 206.9224 | 0.500301 | 0 | 102.364 |
| min | 0.174625 | 12.68 | 24.10099 | 3.20427 | 17.23 | 0 | 0 | 7.431696 |
| 25% | 0.933159 | 18.16414 | 48.4226 | 9.05778 | 78.01 | 0 | 0 | 58.10649 |
| 50% | 1.203991 | 19.755 | 57.6268 | 10.26054 | 149.68 | 1 | 0 | 137.6667 |
| 75% | 1.516648 | 23.84762 | 65.12767 | 12.38454 | 255.36 | 1 | 0 | 216.0659 |
| max | 2.410672 | 29.78656 | 88.55074 | 24.32628 | 1051.83 | 1 | 0 | 554.8244 |

**Supplementary figure 1:** Corraletion chart of the datasets features. (A) Training dataset. This chart is partly presented in Figure 2 And is fully presented here for easy comparison. (B) Holdout dataset. Standard Pearson correlation coefficient is presented.

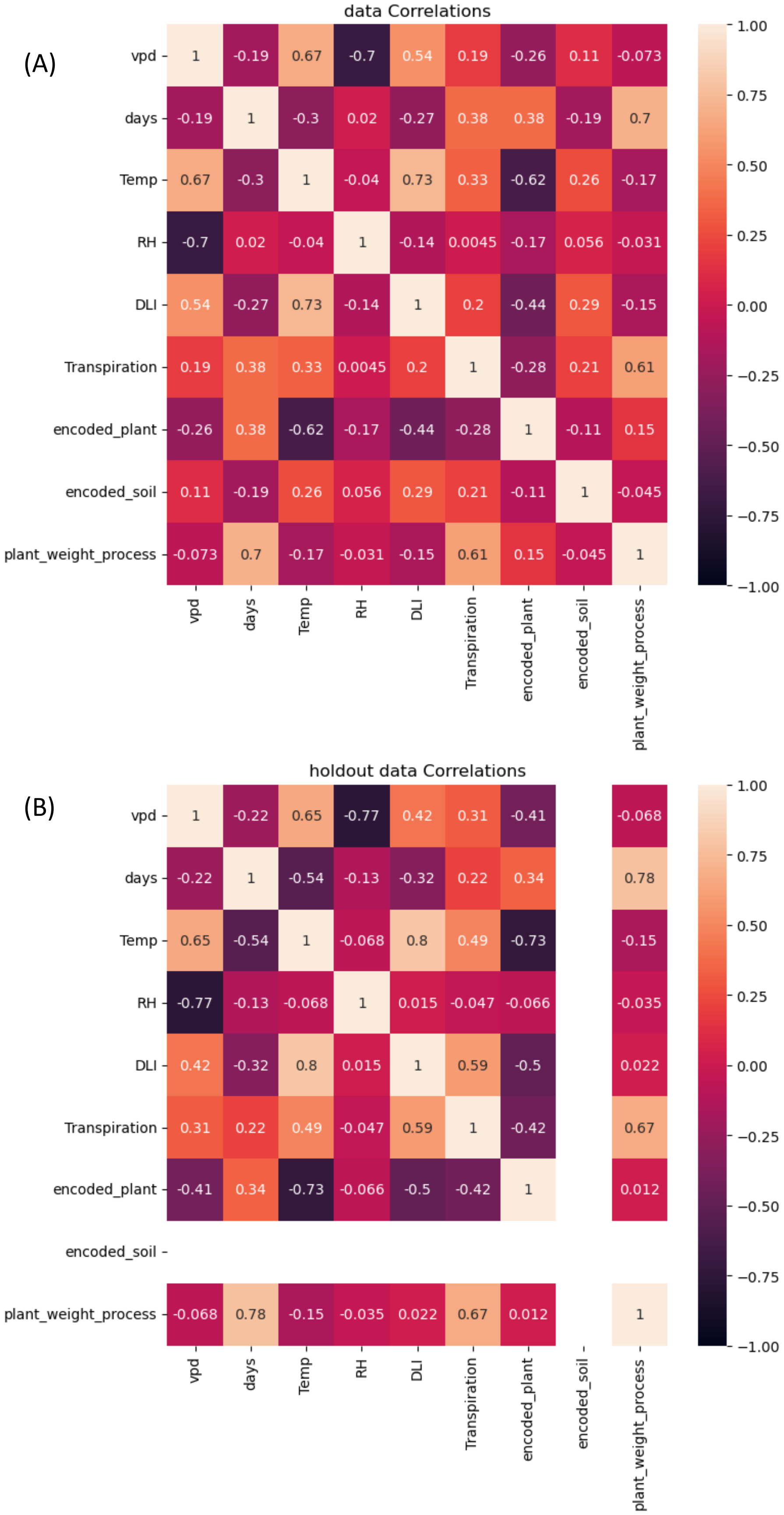

**Supplementary figure 2: Environmental conditions and experimental setup in the indoor growth room (Room 101).** (A) Photograph of the growth room used in the second external dataset, located within the I-CORE center. Tomato plants were grown in individual pots placed on load-cell lysimeters for continuous weight monitoring. (B) Environmental conditions over two consecutive days, illustrating the relatively stable indoor climate. From top to bottom: light intensity (PPFD, µmol $m^{-2}$ $s^{-1}$), relative humidity (%), temperature (°C), and vapor pressure deficit (kPa). These controlled conditions differ from the more variable environment of the main greenhouse, offering a distinct and independent setting for model validation.

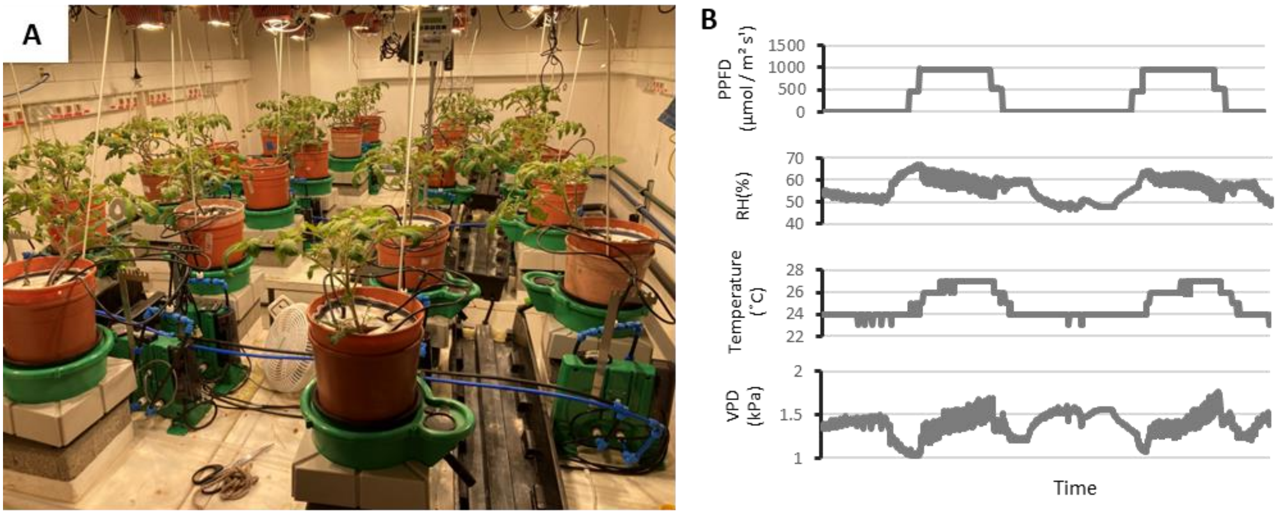

**Supplementary figure 3: Distribution of Environmental and Physiological Parameters by Soil and Crop Type.** This figure presents the distribution of key environmental and physiological parameters stratified by soil type (top row) and crop type (bottom row). The histograms illustrate the frequency distribution of temperature (Temp), relative humidity (RH), vapor pressure deficit (VPD), daily light integral (DLI), plant weight process, and transpiration. Top Row: Comparison by encoded soil type, where 0 represents sand (4630 observations) and 1 represents soil (1607 observations). Bottom Row: Comparison by encoded plant type, where 0 represents tomato (4082 observations) and 1 represents cereal (2155 observations).

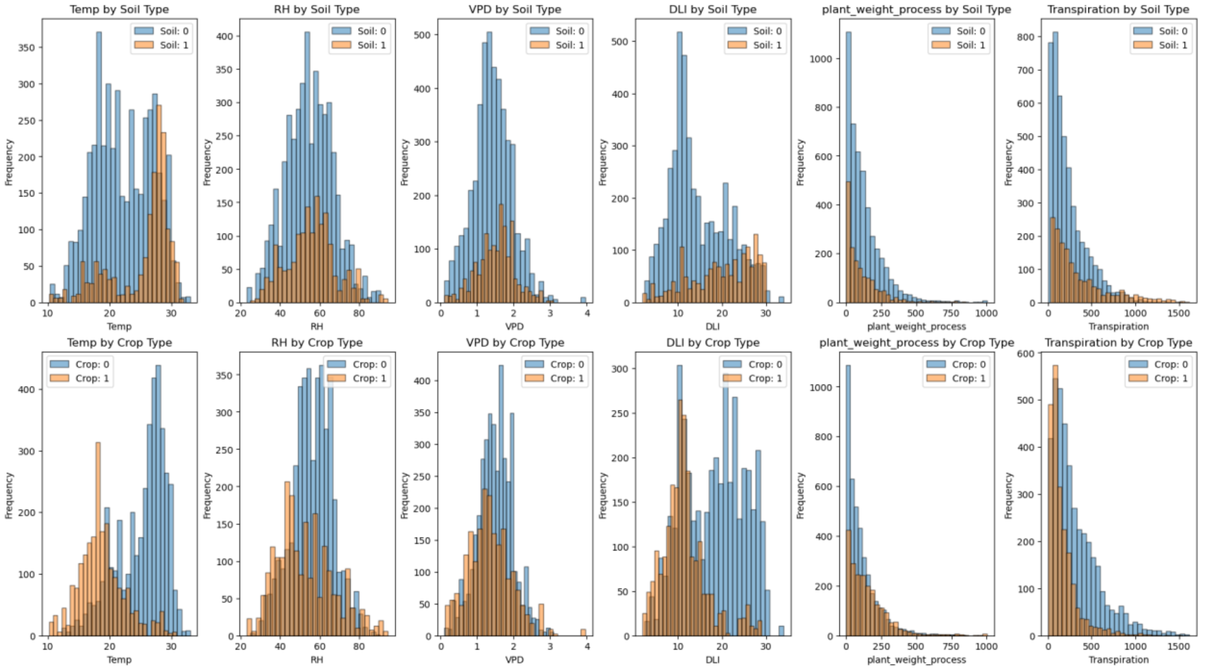

**Supplementary figure 4:** SHAP feature importance. This figure demonstrates two examples of SHAP (SHapley Additive exPlanations) feature importance analysis for predicting plant transpiration, with a base value of 261 grams. The analysis highlights how different features including soil type, plant weight, Relative Humidity (RH), plant type, Daily Light Integral (DLI), VPD, and temperature, significantly influence the prediction in either direction. Reding these plots from the lowest features to the top and looking how etch feature shifts the prediction.  The top graph shows a final prediction of 544.6 grams, where high plant weight is the dominant factor contributing to an increase, alongside encoded soil and RH. Conversely, the bottom graph displays a lower prediction of 152.14 grams, primarily due to a strong negative contribution from the low temperature. Despite the positive influence of plant weight and soil type, the low temperature significantly limits the final prediction, illustrating the complex interplay of factors.

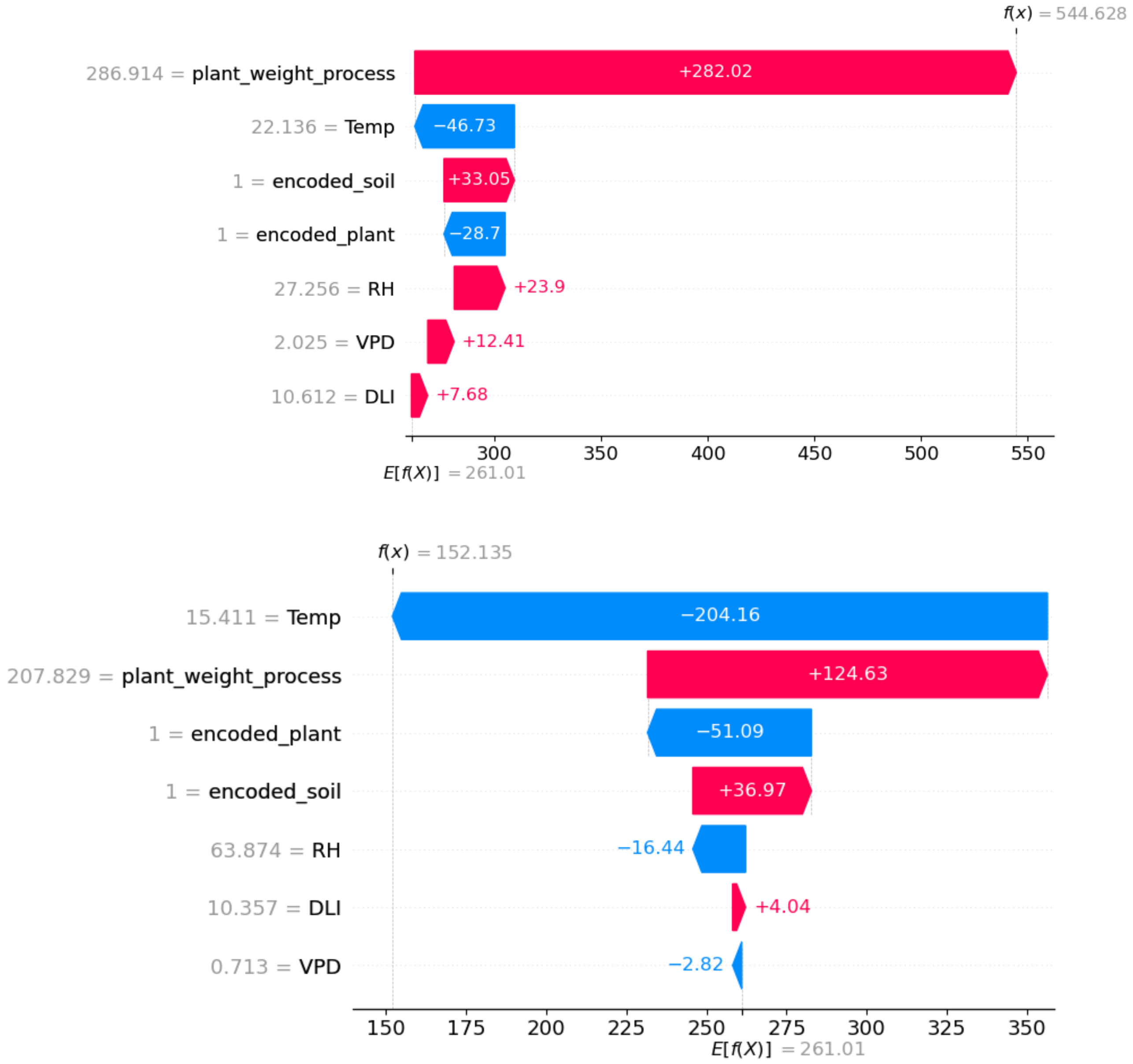

**Supplementary figure 5:** Distribution of transpiration . (A) Box and wisker plot of transpiration in the data used to train the models (left) and in the holdout data (right). T-test significant different

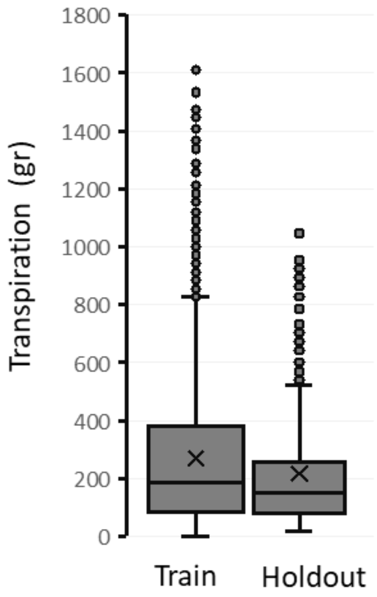

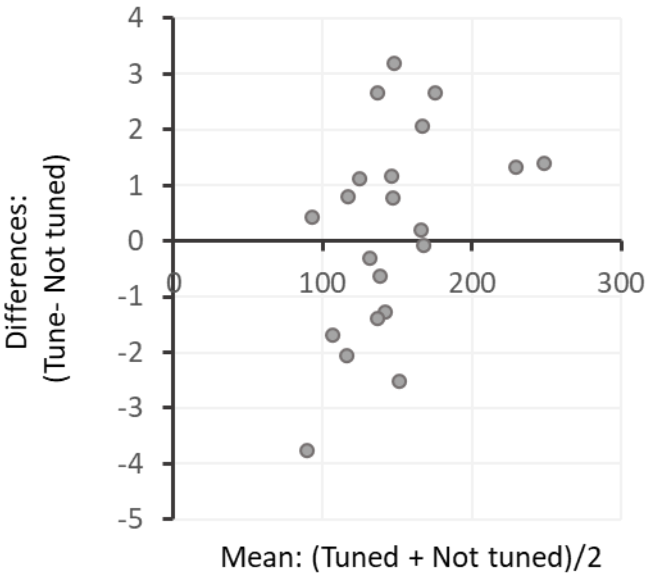


Mean RMSE
Tuned: 146.58
Not tuned: 146.38

**Supplementary figure 6: Model Tuning did not improve Performance.** This figure illustrates the prediction performance of a tuned versus non-tuned random forest model on holdout data. The model was evaluated 21 times, each with a different random seed, resulting in different holdout subsets each time. In each evaluation, the model was tested with both tuned hyperparameters (optimized on 90% of the training data) and default hyperparameters (not tuned). The differences in RMSE scores between the tuned and non-tuned models are depicted. The mean difference in RMSE is 0.19, with a p-value of 0.63, indicating that the tuning process did not improve model performance.
